# Alternative polyadenylation of a polycistronic locus regulates the tumor-suppressive miR-317 and the lncRNA Peony in Drosophila

**DOI:** 10.1101/2024.10.10.617551

**Authors:** Travis D. Carney, Halyna R. Shcherbata

## Abstract

The impact of non-coding RNAs on stem cell biology and differentiation processes is an important and incompletely understood area of research. Using the testes of *Drosophila melanogaster* as a valuable system for investigating these processes, we identified a polycistronic locus from which two non-coding transcripts, *miR-317* and the long non-coding RNA (lncRNA) *Peony*, are produced, with alternative polyadenylation implicated in regulation of their differential expression levels. We report here that each transcript has a distinct role in Drosophila testes; the increased expression of *Peony* results in the disruption of the muscle sheath covering the testis, and the absence of *miR-317* leads to the emergence of germ-cell tumors in developing flies. The deficiency of *miR-317* increases Notch signaling activity in somatic cyst cells, upregulates multiple predicted targets of *miR-317*, and drives germline tumorigenesis. Our findings establish miR-317 as a tumor suppressor controlling Notch signaling strength and uncover alternative polyadenylation as an unrecognized mechanism governing miRNA expression.

## Introduction

More than three decades of research have revealed the involvement of microRNAs (miRNAs) in many aspects of stem cell biology and in processes of differentiation. MiRNAs have been shown to modulate cell division in human embryonic stem cells by targeting key cell cycle checkpoint components (Qi et al. 2009). In Drosophila, miRNAs are required for cell cycle progression as well as maintenance of ovarian germline stem cells (GSCs) (Hatfield et al. 2005; Shcherbata et al. 2007). In addition to roles in stem cell function and homeostasis, we have previously shown that miRNAs affect aspects of cell fate determination in multiple cell types in Drosophila (Kucherenko et al. 2012; Yatsenko and Shcherbata 2014). These studies demonstrate the importance of miRNAs in the crucial processes of stem cell function and cellular specification and differentiation.

The Drosophila testis has proven to be an invaluable model for the study of stem cell biology, intercellular signaling, and differentiation. The testis is a coiled tube, closed at the apical (anterior) end, in which the processes of spermatogenesis take place (Figure 1, A-D). At the testis apex, a cluster of specialized somatic cells called the hub forms a stem cell niche capable of maintaining two populations of stem cells: GSCs and somatic cyst stem cells (CySCs) (de Cuevas and Matunis 2011) (Figure 1, B and D). Both are maintained through direct contact with the hub and through multiple cell-signaling pathways. GSCs and CySCs divide asymmetrically such that one daughter retains both niche contact and stemness, while the other daughter, no longer proximal to the self-renewal-promoting signals from the niche, begins a differentiation program. The differentiating daughter of the GSC, a gonialblast, becomes encapsulated by two somatic daughter cells, termed cyst cells, which themselves differentiate in concert with the germ cells and maintain this association throughout spermatogenesis. Cyst cells serve as a differentiation niche, supporting spermatogenesis processes and protecting germ cells from exogenous signals (Decotto and Spradling 2005; Zoller and Schulz 2012; Papagiannouli 2014). The gonialblast undergoes four rounds of transit-amplifying mitotic divisions to form a 16-cell cluster of cells termed spermatogonia (Greenspan et al. 2015). This cell cluster and the two encapsulating cyst cells compose a single cyst. Spermatogonial cells mature into primary spermatocytes, substantially increase their volume, and undergo meiosis to form spermatids, each of which elongates, matures, and is released as a mature sperm to be stored in the seminal vesicle until copulation. Therefore, a single GSC division has the potential to result in the formation of 64 haploid sperm (Greenspan et al. 2015).

**Figure 1.**
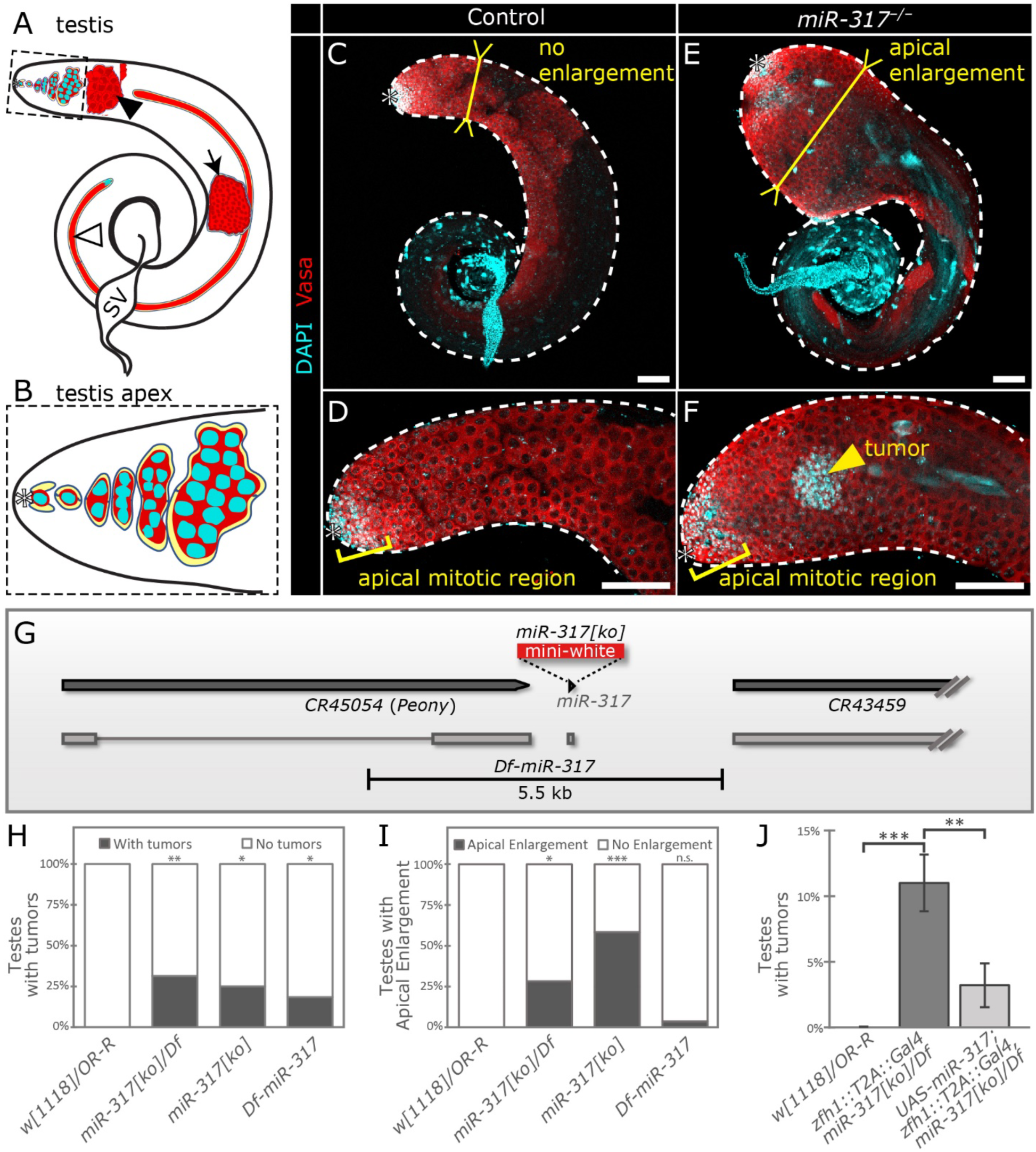
*miR-317* mutation causes testis phenotypes in adult males. **A** Schematic of one adult testis. Shown are stem cell niche (hub; denoted by asterisk); somatic cyst cells (yellow) surrounding germline cysts; 16-cell spermatocyte cyst (solid arrowhead); 64-cell post-meiotic cyst (black arrow); elongated spermatid-stage cyst (open arrowhead); and seminal vesicle (SV). Dashed box outlines the apex, enlarged in B. **B** Testis apex. Hub (asterisk); germline cells (red) surrounded by somatic cyst cells (yellow). Germline cells divide to form 16-cell cysts. **C** Control testis is a coiled, closed-ended tube (outlined with white dashed line). The apical half of the testis has a roughly constant wideness (indicated by yellow bar), until tapered & rounded at the apex. **D** Control testis apex. **E** *miR-317^-/-^*testis exhibiting apical enlargement, a dramatic widening of the testis tube near the apical end, indicated by yellow bar. **F** *miR-317^-/-^*apex containing a tumor (yellow arrowhead), identified by strong expression of Vasa (red) as well as strong DAPI staining (cyan) removed from the apical mitotic region at the apex. **G** Genomic context of *miR-317*, which is closely apposed to the lncRNA gene *CR45054* (*Peony*). *miR-317[ko]* is a knock-in mutant in which the pre-miRNA is replaced by a *mini-white* cassette. *Df-miR-317* is a 5.5-kb deletion of *miR-317* and the large second exon of *Peony*. **H-I** Quantification of the percent of testes exhibiting tumors (H) or apical enlargement (I). **J E** Quantification of percent of testes bearing tumors in a rescue experiment using *zfh1::T2A::Gal4* to express *miR-317* in a mutant background. Controls exhibit no tumors. Overexpression of *miR-317* significantly rescues the mutant’s tumor formation phenotype. In E, error bars indicate standard deviation from the mean; Student’s unpaired t-test was utilized to test for statistical difference between genotypes. In H and I, chi-square analysis was used to determine statistical significance; n=22-35 testes per genotype. *, p<.05, **, p<.01; ***, p<.001. Scale bars in C-F are 100 µm.

The processes of proliferation and differentiation must be carefully balanced to ensure that each cyst produces the proper number of gametes while avoiding tumor-like overgrowth. In particular, precisely four spermatogonial mitotic divisions must occur to form a normal, 16-cell spermatocyte cyst. Numerous instances have been reported in which these processes fail and result in overproliferation. The genes *bag of marbles* (*bam*), *benign gonial cell neoplasm* (*bgcn*), and *Rbfox1* act cell-autonomously in the germline, and mutation of these genes results in overproliferation of early, undifferentiated germ cells (McKearin and Spradling 1990; Gonczy et al. 1997; Tastan et al. 2010). In addition, genes affecting signaling, transport, and numerous other processes in the somatic cyst cells have been shown to affect the germline non-autonomously, interfering with the differentiation program and causing dramatic overproliferation of undifferentiated germ cells (Matunis et al. 1997; Kiger et al. 2000; Tran et al. 2000; Joti et al. 2011; Fairchild et al. 2017; Tang et al. 2017). From these studies, a clear picture has emerged in which the close association and signaling relationship between germline and somatic cells is crucial for proper spermatogenesis.

Recent studies have shown that lncRNAs are expressed in a tissue-specific manner and are particularly enriched in the testes of Drosophila and other animals, including mammals (Cabili et al. 2011; Young et al. 2012; Brown et al. 2014; Necsulea et al. 2014). Many testis-expressed lncRNAs are enriched in late developmental stages, including meiotic and elongated cysts (Wen et al. 2016; Vedelek et al. 2018). Some of them have roles in the late stages of spermatogenesis, as mutations cause spermatid defects including defective chromatin compaction and sperm individualization (Wen et al. 2016).

We are interested in elucidating the effects of non-coding RNAs (ncRNAs), in particular miRNAs, on processes related to stem cell biology and differentiation and how these processes can be perturbed when miRNAs are dysregulated. Also important is the identification of miRNA target genes that are in turn affected in these mutants. To this end, we screened a number of miRNAs reported to exhibit expression in testes and/or ovaries. We closely examined miRNA-mutant gonads for phenotypes consistent with stem cell division/self-renewal problems or differentiation defects.

Here we report that deficiency of the miRNA *miR-317* causes the formation of germ-cell tumors in developing fly testes. Previous reports have implicated *miR-317* in several processes and behaviors, including negative regulation of the Toll immune response (Li et al. 2017; Li et al. 2019); larval ovary morphogenesis and development (Yang et al. 2016); startle-induced locomotion (Yamamoto et al. 2008); aggressive behavioral response (Edwards et al. 2009); female response to mating (Fricke et al. 2014); and embryonic regulation of Cyclin B expression (Pushpavalli et al. 2014). Furthermore, overexpression (OE) of *miR-317* is sufficient to prevent the formation of imaginal disc tumors in the potently oncogenic background of simultaneous *RasV12* OE and *discs large* mutation (Wang et al. 2022). However, the present study is the first to demonstrate an endogenous role of *miR-317* as a tumor suppressor.

The *miR-317* mutant allele *miR-317[ko]* results in the concomitant upregulation of a co-transcribed lncRNA *CR45054* (FBgn0266414), which causes a disruption of the muscle sheath surrounding the testis and a pronounced enlargement of the testis apex. As this distinct phenotype causes the testis to resemble the appearance of a peony flower, we have named the lncRNA *CR45054* as “*Peony.*” Our data indicate that *miR-317* and *Peony* are mutually expressed from a single polycistronic non-coding RNA locus and that their expression depends on differential polyadenylation.

Furthermore, we find that somatic OE of either of two putative *miR-317* targets, Rotund (Rn) or CG18265, or of the Notch signaling cofactor, Mastermind (Mam), is sufficient to cause the formation of testis tumors. All three are transcriptional regulators. Deficiency of *miR-317* results in increased Notch pathway signaling activity in somatic cyst cells, and the gene dose reduction of *mam* results in a partial abrogation of tumor formation, potentially implicating the Notch pathway in this phenotype. Our results show that *miR-317* is a novel tumor suppressor, and this important function requires the repression of multiple targets in the somatic cells of the Drosophila testis.

## Results

### *miR-317[ko]* males have multiple testis phenotypes

The Drosophila testis serves as a convenient system for detecting defects in division and maintenance of stem cells as well as the differentiation of their progeny. In searching for miRNA genes involved in these processes, we examined the gonads of a loss-of-function (LOF) mutant, *miR-317[ko]*, in which the pre-miRNA has been precisely replaced with a *mini-white* cassette by ends-out homologous recombination (Chen et al. 2014) (Figure 1, E-G). In contrast to controls, in which the apical half of the testis maintains a roughly uniform width until the rounded apex (Figure 1C), we found that some *miR-317[ko]* testes exhibit a dramatic enlargement at the testis apex (apical enlargement, AE; Figure 1E). At the apex of control testes, the mitotic region containing transit-amplifying spermatogonia is characterized by bright DAPI staining, owing to small cell size and tightly condensed chromatin. As germ cells mature into spermatocytes, the cells grow substantially and their chromatin decondenses, leading to much more diffuse DAPI staining (Figure 1, C and D). In addition to the AE phenotype, we also observed an incidence of tumors in *miR-317[ko]* testes, identifiable by very bright DAPI staining that is apart from the apical mitotic region, as well as the expression of Vasa, an RNA helicase commonly used as a germ-cell marker (Figure 1F). Interestingly, these tumors are found to exist either with or without the AE phenotype.

To reduce the likelihood of genetic background-induced phenotypic artifacts, we generated a new deficiency allele using FLP-mediated recombination between FRT sites present in two transposon insertions flanking *miR-317* (Parks et al. 2004). This resulted in the deletion of 5.5 kb that removes *miR-317* as well as the large second exon of the nearby lncRNA gene, *Peony* (Figure 1G). We predict that this deletion, which we call *Df-miR-317*, represents a LOF allele for both *miR-317* and *Peony*.

We compared testes from homozygous (*miR-317[ko]*) and hemizygous (*miR-317[ko]/Df*) adult males and found that testis tumors exist in both genotypes with similar frequency, strongly suggesting that this phenotype is attributable to the loss of the miRNA (Figure 1H). Interestingly, we noted that while *miR-317[ko]* and *miR-317[ko]/Df* testes exhibit the AE phenotype, we very rarely observe this phenotype in *Df-miR-317* testes (Figure 1I). As the primary difference between *miR-317[ko]* and *Df-miR-317* is the deletion of *Peony* only in the latter, this discrepancy raises the possibility that the lncRNA is involved in the development of the AE phenotype. In all subsequent experiments requiring *miR-317* LOF, we use either *miR-317[ko]* homozygotes or *miR-317[ko]/Df* flies.

To investigate the role of *mir-317* in tumor formation, we performed a rescue experiment using an early cyst-cell driver, *zfh1::T2A::Gal4*, in which the self-cleaving peptide T2A and Gal4 sequences have been fused to the C-terminus of Zfh1 by CRISPR-Cas9-mediated genome editing (Albert et al. 2018). We found that *miR-317[ko]/Df* combined with a heterozygous *zfh1::T2A::Gal4* allele results in an incidence of tumors of about 11%, and addition of *UAS-miR-317* elicits a significant rescue, reducing the incidence of tumors to about 3% (Figure 1J). This result strongly suggests that *miR-317*’s tumor suppressive activity takes place in the early somatic cells.

Based on these analyses, we hypothesize that, although encoded by the same genomic locus, these transcripts have distinct roles — the *Peony* lncRNA in testis morphogenesis and *miR-317* as a tumor suppressor.

### RNA-seq in *miR-317* mutant testes reveals differential expression of genes important for cell signaling and differentiation

To attain a global view of the genes perturbed in *miR-317* mutants, we performed RNA-seq analysis comparing testes from a control genotype, *w[1118]*, to the LOF mutant, *miR-317[ko]* (see Materials and Methods). We detected 13232 total genes in testes, 1844 (13.9%) of which are dysregulated (Figure 2A; Supplemental Table 1). We performed a Gene Ontology (GO)-term enrichment analysis to investigate the biological processes that are overrepresented among the dysregulated genes. Enriched terms include signal transduction and cell differentiation, as well as cell communication, cell migration, and morphogenesis, all of which processes are relevant to tumorigenesis like that we discovered in testes (Figure 2B).

**Figure 2.**
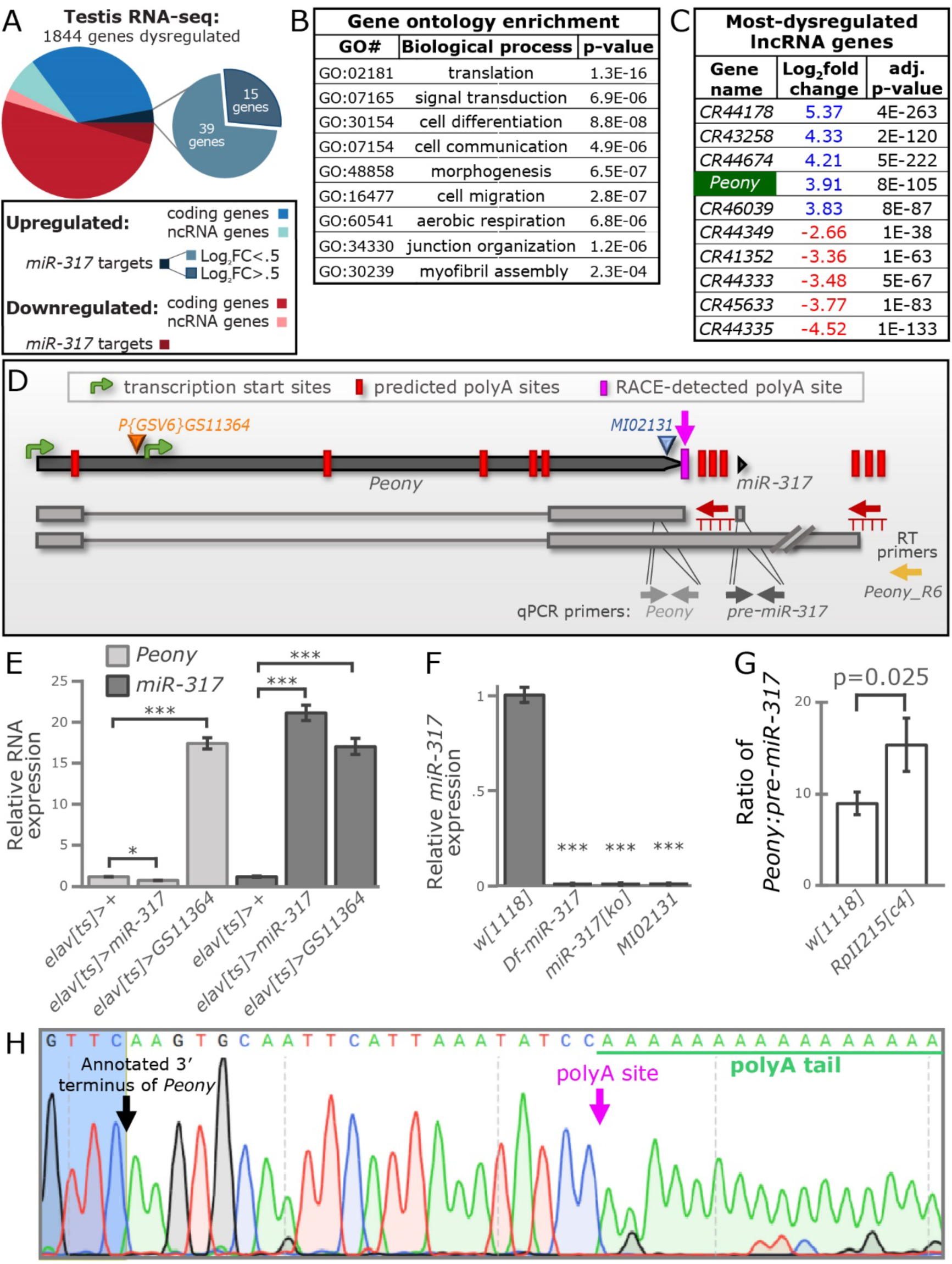
*Peony* is upregulated in *miR-317[ko]* and affects testis morphology. **A** Results of RNA-seq experiment comparing control (*w[1118]*) and mutant (*miR-317[ko]*) testes. Downregulated genes are indicated in shades of red; upregulated genes in blues; non-coding genes in light blue and pink. *In-silico-*predicted *miR-317* targets are shown (dark blue and dark red); those predicted targets upregulated in mutant testes were analyzed further regarding the magnitude of upregulation (Log_2_FC>0.5 or Log_2_FC<0.5; blue-gray pie chart). **B** Gene ontology (GO) analysis of the dysregulated genes; several of the most enriched ‘biological process’ GO terms are shown. For each process shown, the corresponding GO number is given as well as the significance of enrichment. **C** The five most upregulated and downregulated lncRNAs are shown. The Log_2_FC values are shown in blue if upregulated, red if downregulated, as in Supplemental Tables. *Peony* is one of the most upregulated lncRNAs in *miR-317* mutants (highlighted in green). **D** Schematic of the *Peony-miR-317* locus showing predicted transcription start sites and polyA sites; polyA site detected using 3′RACE; transgenic insertions used in this study; and locations of primers used for reverse transcription and qPCR. The most distal polyadenylation sites are not annotated and are predicted based on this work. Transcripts (light gray rectangles) depict putative alternative isoforms that differ based on alternative polyadenylation. **E** Results of qRT-PCR using qPCR primers in *Peony* and pre-*miR-317.* Control expression levels were compared with overexpression using *elav-Gal4; Tub-Gal80[ts]* (*elav[ts]*). **F** Mature *miR-317* levels in control (*w[1118]*) and three mutant genotypes: *Df-miR-317, miR-317[ko],* and MiMIC transposon *MI02131*. **G** *Peony:*pre*-miR-317* ratios resulting from qRT-PCR from two genotypes: *w[1118]* and *RpII215[c4]* using the qPCR primers shown in panel D. Shown are mean ± standard deviation from three biological replicates; statistical significance determined using a Student’s t-test. **H** Sequencing result of 3′RACE shows the site of cleavage and polyadenylation (polyA site) 22 bp downstream of the annotated 3′ end of *Peony*. Graphs show mean ± standard deviation. Statistical significance was assessed using Student’s t-test based on technical triplicates (E&F) or biological triplicates (G). *, p<.05; ***, p<.001.

Consistent with previous reports (Djebali et al. 2012; Brown et al. 2014; Wen et al. 2016; Joshi and Rajender 2020; Hong et al. 2021), we detected a substantial proportion of lncRNA genes in testes (Supplemental Table 1; Figure 2A). In contrast to protein-coding genes, lncRNAs are more likely to be up-than downregulated; indeed, the five most-upregulated genes, and more than a third of the top 100 most-upregulated genes, are lncRNAs (Supplemental Table 1). Interestingly, among those lncRNA genes strongly upregulated in *miR-317[ko]* testes is *Peony,* the gene just upstream of *miR-317* (Figure 2C).

### Upregulation of *Peony* causes disruption of the testis muscle sheath

Analyses of mutant testes revealed that AE is concomitant with a disruption of the muscle sheath that surrounds the testis (Supplemental Figure 2, A-C). While nearly 80% of *miR-317[ko]* testes exhibit some degree of muscle disruption, *Df-miR-317* testes muscles appear intact as in controls (Supplemental Figure 2D). Since *Peony* is strongly upregulated in *miR-317[ko]* (Supplemental Table 1; Figure 2C), we generated a *UAS-Peony* transgene to allow conditional OE of a full-length cDNA. To determine whether *Peony* plays a role in the AE or muscle disruption phenotype, we overexpressed *Peony* using Gal4 lines that express in various cell types in testes (Supplemental Figure 1). Indeed, we found that muscle disruption is evident upon *Peony* OE in some cell types. Expression of *Peony* in somatic cyst cells or germ cells – using *tj-Gal4* or *vasa-Gal4*+*nos-Gal4*, respectively – causes muscle disruption in 13 or 34% of testes, respectively (Supplemental Figure 2D; Supplemental Figure 1, A and B). OE using *bab1-Gal4*, expressed in somatic cyst cells, muscle cells, and pigment cells (Supplemental Figure 1, C and D), causes muscle sheath disruption in 42% of testes. OE with *Mef2-Gal4*, expressed in muscle cells, causes muscle sheath disruption in 33% of testes, while OE with *MJ12a-Gal4*, which drives expression only in the pigment cell layer that also surrounds the testis (Hrdlicka et al. 2002) (Supplemental Figure 1, E and F), does not cause muscle disruption (Supplemental Figure 2D).

Since none of the conditions described above results in a muscle disruption phenotype as severe as *miR-317[ko]*, we turned to ubiquitous OE using *Tub-Gal4*. This condition causes a high degree of lethality, so we used a *Tub-Gal4; Tub-Gal80[ts]* combination (*Tub-Gal4[ts]*) to block expression until late larval stages. This results in a strong incidence of muscle disruption very similar to that seen in *miR-317[ko]* testes (Supplemental Figure 2D). Moreover, while OE using the other cell-type-specific drivers fails to result in AE, this phenotype does manifest upon ubiquitous OE (Supplemental Figure 2, E and F). Increased *Peony* levels do not, however, result in the formation of tumors with any driver. We conclude that while tumor formation is due to a loss of *miR-317* function, the AE phenotype is a result of the upregulation of *Peony* that occurs in *miR-317[ko]* testes or upon exogenous OE.

### Differential levels of *Peony* and *miR-317* rely on alternative polyadenylation

We next investigated the relationship between *Peony* in *miR-317* expression. It has been proposed that *Peony* serves as the primary miRNA transcript from which *miR-317* is processed (Kadener et al. 2009). We confirmed this first by driving expression from a UAS insertion in the intron of *Peony*, *P{GSV6}GS11364* (Figure 2D), which is sufficient to cause upregulation of *miR-317* (Figure 2E). Next, we assayed *miR-317* levels in flies homozygous for *MI02131,* a MiMIC gene trap containing a transcriptional stop signal (Venken et al. 2011) (Figure 2D). Similar to *miR317[ko]* and *Df-miR-317* flies, *MI02131* causes a strong loss of *miR-317* expression (Figure 2F). These results suggest that *Peony* expression is required and sufficient for production of *miR-317*.

In silico analysis (RegRNA 2.0) (Chang et al. 2013) predicts three polyadenylation (polyA) sites between *Peony* and *miR-317* (Figure 2D), raising the interesting possibility that the differential usage of polyA sites may play a role in regulating the expression of *miR-317*. To investigate this possibility, we extracted RNA from control testes, completed reverse transcription using an oligo-dT primer, and performed qPCR using two sets of primers: one targeting the pre-miRNA region and another in *Peony* (Figure 2D). In this way, we were able to quantify both regions of the parent transcript and calculate a *Peony:miR-317* ratio. This ratio is 9:1 in control testes (Figure 2G), suggesting that *Peony* is greater in abundance than *miR-317*. To test the hypothesis that this difference is due to alternative polyA sites, a proximal site between the two genes and a distal site downstream of the miRNA, we made use of *RpII215[c4],* an RNA polymerase II subunit mutation that results in reduced transcription elongation speed. This slowed elongation imparts to the transcription apparatus the tendency to utilize a proximal polyA site (Liu et al. 2017), in effect forcing a proximal polyA-site bias. We found that compared to controls, the *Peony:miR-317* ratio increases nearly twofold in *RpII215[c4]* testes, from 8.5 to 15.3 (p=.025, Figure 2G). This result is consistent with the differential utilization of a polyA site upstream of *miR-317*. Thus, we used a modified 3′ rapid amplification of cDNA ends (3′ RACE) approach (see Materials and Methods) and confirmed the presence of a cleavage and polyA site 22 bp downstream of *Peony*’s annotated 3′ end (Figure 2H). These results strongly suggest that, in testes, the relative levels of *Peony* and *miR-317* are affected by the differential choice of either a proximal or a distal polyA site.

### *miR-317* LOF tumors consist of GSC-like and early spermatogonial cells

Next, we turned our attention from the lncRNA, *Peony*, to examine the tumor suppressive functions of *miR-317*. To more fully characterize the *miR-317[ko]* testis tumor phenotype, we stained control and mutant testes with antibodies that mark different cell types and indicate different stages of differentiation. The strong expression of Vasa and Adducin (Add) suggests that the tumors are largely composed of germ cells, not somatic cells (Figure 3, A and A′). We used several additional means to verify this. Add interacts with the actin cytoskeleton and labels distinct germline-specific structures in early germ cells: spectrosomes and fusomes. The morphology of these structures can indicate the identity (i.e., differentiation state) of these cells: spherical spectrosomes are indicative of a GSC or gonialblast (the immediate daughter of a GSC) identity, while branched fusomes indicate a more mature spermatogonial identity. We found that the tumors in *miR-317* mutants possess mostly spectrosomes but can also contain a smaller amount of branched fusomes, sometimes within a single tumor (Figure 3A′′). These results demonstrate that the majority of the cells within the tumors have an early germ-cell identity.

**Figure 3.**
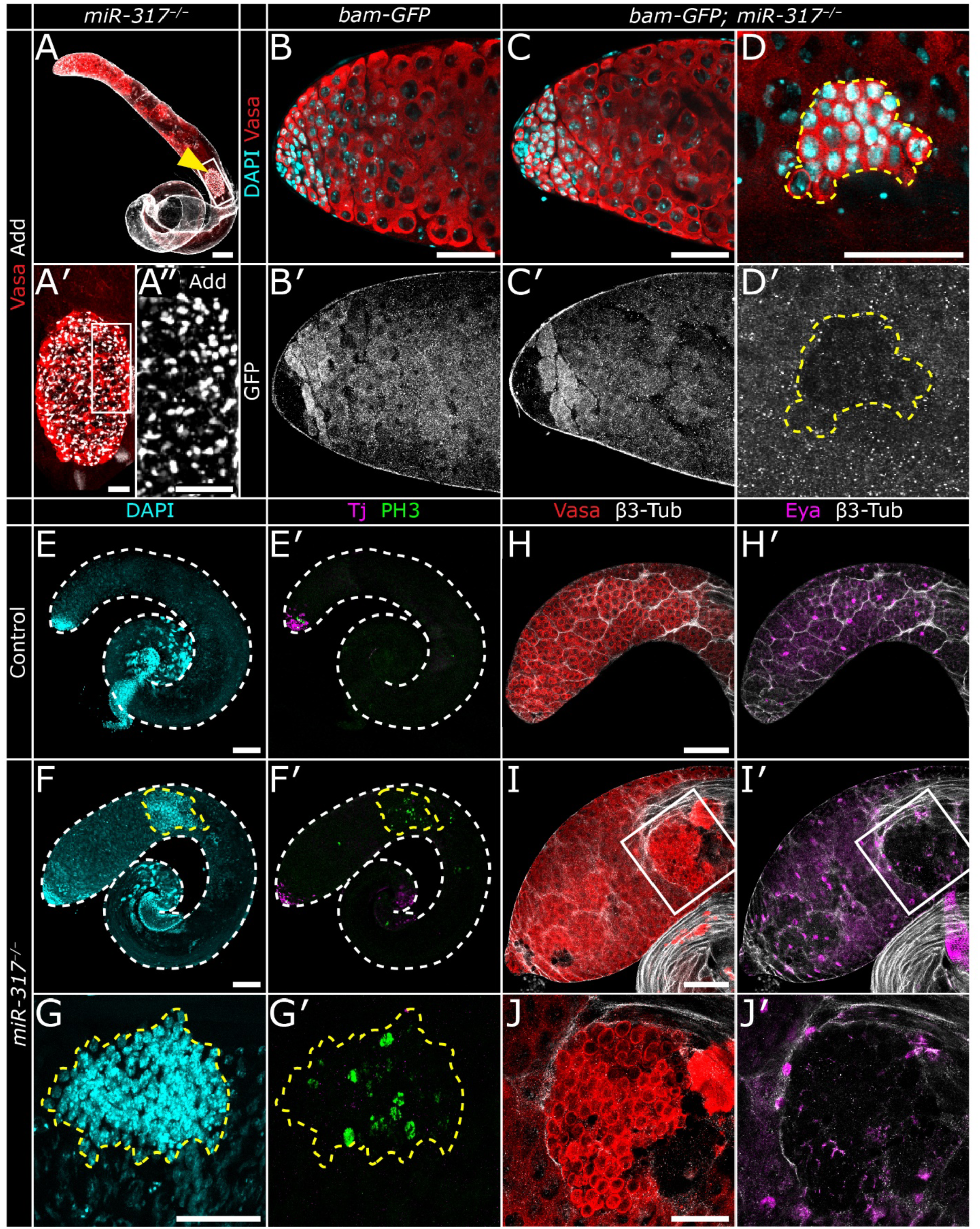
*miR-317* mutant testis tumors are composed of early germline cells. **A** *miR-317^-/-^* testis exhibiting a tumor (yellow arrowhead) with strong Vasa (red) and Adducin (Add; grayscale) staining. **A′** Enlargement of the tumor outlined with a white rectangle in A. All cells are Vasa-positive and Add-positive. **A′′** Enlargement of the white rectangle in A′ shows Add staining exists mostly in spherical-shaped spectrosomes, with a minority exhibiting minimal branching. **B** Apex of testis with a *bam* transcriptional reporter (*bam-GFP*). GFP is shown in single panel in B′. **C** *bam-GFP* in a *miR-317^-/-^*background; GFP pattern (**C′**) does not appear different from control (**B′**). **D** Tumor in a *bam-GFP; miR-317^-/-^* testis; tumors of this genotype do not express *bam-GFP* (**D′**). **E** Control testis (outlined with white dashed line) exhibiting strong DAPI staining near the apex, in the region where germline cells undergo mitotic divisions. **E**′ Tj-positive cyst cells reside near the testis apex, in proximity of the mitotic zone. PH3-positive cells mark mitotically active germline cells, which divide in synchrony within each germline cyst. **F** *miR-317^-/-^* testis with a large tumor, outlined by yellow dashed line and identified by region of bright DAPI-positive cells removed from the apex. **F**′ Same testis as shown in F, now showing that while some cells in the tumor are PH3-positive, the Tj-positive cells are present only near the apex as in controls. **G** A *miR-317^-/-^* tumor with bright DAPI staining. **G**′ Same tumor as shown in G. Some scattered cells are PH3-positive; tumor and surrounding cells are Tj-negative. (E,F, & G: DAPI, cyan; PH3, green; Tj, magenta). **H** Apex of control testis. Vasa marks germline cells, and β3-Tub marks cyst cell cortexes. **H**′ Same testis as in H; Eya marks somatic cyst cell nuclei. **I** *miR-317^-/-^* testis stained with Vasa and β3-Tub and exhibiting a tumor (indicated by white box). **I**′ Same testis as in I; β3-Tub and Eya mark cyst cell cortexes and nuclei, respectively. **J-J**′ Enlargement of the tumor shown in I & I′ showing that while Vasa is strongly expressed in the tumor, β3-Tub and Eya are excluded. (H-J: Vasa, red; β3-Tub, grayscale; Eya, magenta). A, E, F, H, I: scale bars 100 µm. B, C, D, G, & J: scale bars 50 µm. A′, A′′: scale bars 20 µm.

The pro-differentiation protein Bam is an indicator of the differentiation state of germ cells. In control testes, Bam is expressed in germ cells of 4- to 8-cell cysts and is absent from GSCs, gonialblasts, and 2-cell cysts. *bam-GFP* is a reporter construct consisting of *bam* regulatory elements driving the expression of GFP (Chen and McKearin 2003). As a result, GFP is expressed in the endogenous *bam* pattern: excluded from the most apical germ cells but expressed in 4- and 8-cell cysts. Unlike Bam protein, GFP perdures into later spermatogonial and spermatocyte stages (Figure 3, B and B′). Deficiency of *miR-317* does not change this pattern of GFP expression (Figure 3, C and C′). We found that *bam-GFP* is not expressed in *miR-317[ko]* tumors (Figure 3, D and D′), indicating that in terms of differentiation state, the germ cells comprising the tumor are earlier than the spermatogonia of 4-cell cysts. Taken together, our data suggest that *miR-317* testis tumors consist of early, undifferentiated germ cells similar to GSCs, gonialblasts, or 2-cell cysts. Next, we wanted to determine whether somatic cells also contribute to the cellular composition of the tumors.

The soma-expressed transcription factor, Traffic jam (Tj), is expressed in CySCs and early cyst cells near the testis apex in control and *miR-317* mutants but is not found in cells in or around the tumors in *miR-317* LOF testes (Figure 3, E′-G′). However, we observed that while Tj-positive nuclei cluster close to the apex in controls, in *miR-317* LOF testes, they are found significantly further removed from the apex (Supplemental Figure 3, A-C). This suggests that early cyst cells may exhibit either a delay in differentiation or increased mobility in *miR-317* mutants. The testis hub comprises another population of somatic cells at the apex. We stained these cells with an antibody against Fasciclin III, which allowed us to quantify the number of hub cells. We found that *miR-317[ko]/Df* hubs contain significantly more cells than controls (Supplemental Figure 3D). This further suggests a signaling defect in the somatic cells of the mutant testis apex.

In control testes, GSCs and gonialblasts divide as single cells, then the spermatogonial cells within each cyst divide synchronously in clusters of 2, 4, or 8 cells. After migrating about halfway down the testis, each cyst again divides meiotically in clusters of 16, then 32 cells. As a result, individual, asynchronously dividing germline cells (GSCs and gonialblasts) are normally only found at the apex. To investigate whether *miR-317* LOF tumor cells are proliferative and synchronous, we stained for the mitotic marker, phosphorylated histone H3 (PH3). We found that while some cells in each tumor are PH3-positive, demonstrating mitotic activity, most of the dividing cells are not clustered or synchronous (Figure 3, F′ and G′). This indicates that tumors consist of cells that divide similarly to GSCs or early spermatogonia.

We also stained testes with multiple other somatic cell markers. Eyes absent (Eya) is a transcription factor expressed in mature somatic cyst cells, while β-3-Tubulin marks the cortexes of somatic cells and thus nicely outlines differentiating cysts (Figure 3, H and H′). We found that Eya-positive somatic cells can be found adjacent to tumors, but not within them; similarly, tumors express β-3-Tubulin at their periphery, but this somatic marker does not extend within the tumor (Figure 3, I-J′). Our results lead us to conclude that *miR-317* mutant testis tumors consist not of soma but of germ cells, probably a mix of cells similar to GSCs and early spermatogonia. Hereafter we refer to these tumors as germ-cell tumors (GCTs).

### GCTs arise developmentally and disintegrate in aging testes

As our investigations of the GCT phenotypes described above were limited to young, 0-3-day-old males, we next sought to determine whether the GCT phenotype persists in aging flies. We dissected 1-3-day-old, 2-week-old, and 4-week-old males. Control males do not exhibit GCTs (Figure 4, A, C, and E). We found that approximately the same proportion of mutant testes exhibit GCTs at the 1-3-day-old and 2-week-old time points; however, in older, 4-week-old testes, we found a pronounced decrease in the incidence of GCTs (Figure 4, F and G). These results indicate that new tumors do not form in aging, adult flies.

**Figure 4.**
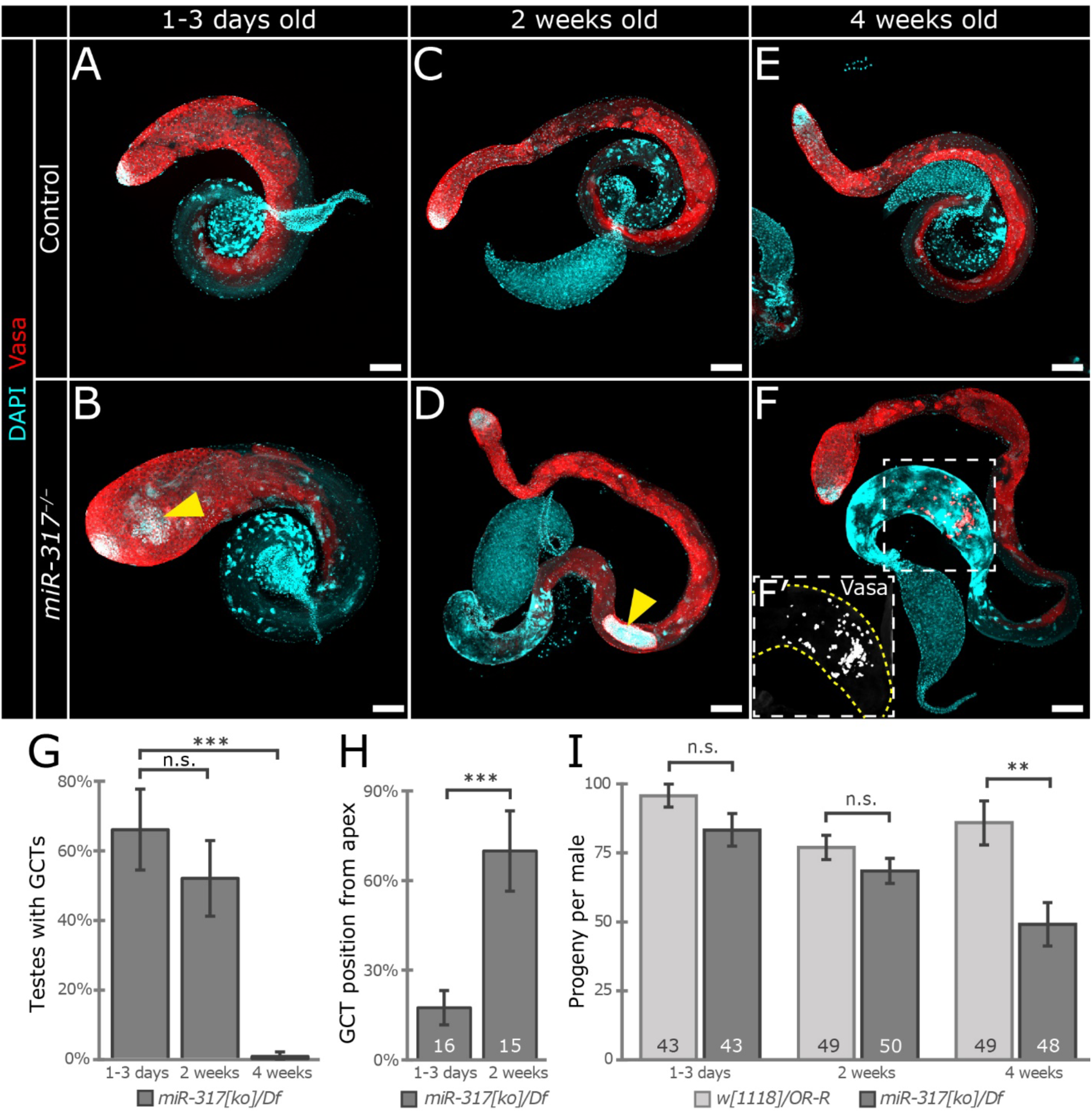
GCTs arise near the testis apex and disintegrate basally in aging testes. **A** Young, 1-3-day-old control testis stained for DAPI (cyan) and Vasa (red). **B** Young *miR-317^-/-^* testis exhibiting a GCT near the apex (yellow arrowhead) with strong Vasa and DAPI staining. **C** 2-week-old control testis. **D** 2-week-old *miR-317* mutant testis with a large GCT in the basal half of the testis (yellow arrowhead). **E** 4-week-old control testis. **F** 4-week-old *miR-317* mutant testis with the partially disintegrated remains of a GCT near the base of the testis (white square with dashed lines), discernible by strong Vasa staining. **F**′ Enlargement of the box in F showing Vasa in grayscale. **G** Graph showing the incidence of GCTs in *miR-317^-/-^* testes over time. **H** Quantification of the average position of GCTs as a percentage from apex to testis base. **I** Fertility results reported as number of progeny produced per male for each genotype and time point. Error bars indicate standard deviation in G-H and SEM in I. Data in G based on 3 or 4 biological replicates. Sample sizes (given as numbers on each bar) indicate number of GCTs counted for each genotype (H) and number of males assayed (I). Student’s t-test used to determine statistical significance. *, p<0.05; **, p<0.01; ***, p<0.001; n.s. = not significantly different. Scale bars 100 µm.

In young *miR-317[ko]/Df* males, as described above, we found GCTs mostly near the apexes of testes (Figure 4B), while in 2-week-old testes, GCTs are located more basally (Figure 4D). We measured the location of GCTs at these time points as a percent displacement between the apex (0%) and the base of the testis, before the seminal vesicle (100%) and found that GCT distance from the apex increases markedly from 17.7% in 1-3-day-old to 70.2% in 2-week-old mutants (Figure 4H). In addition, although very few GCTs exist in 4-week-old mutants, we found evidence of the fragmented remains of tumors near the testis base, in the terminal epithelial region just apical to the seminal vesicles, evidenced by clusters of small cells stained very strongly with Vasa (Figure 4F and F′). 29% of 4-week-old mutant testes exhibit this phenotype, which is never seen in control testes (Figure 4E). We conclude that GCTs in *miR-317* LOF testes originate from germ cells near the apex and then migrate posteriorly through the testis, finally disintegrating in the terminal epithelial region near the testis base. Furthermore, we find that the formation of these GCTs is a phenomenon that occurs during development and in very young flies but does not persist in aging, adult males.

Next, we tested whether *miR-317* deficiency affects male fertility. We performed fertility assays on young, 2-week-old, and 4-week-old males as previously described (Herrera et al. 2021). A subset of both control and mutant males exhibit sterility at all ages. This number is not significantly different in 1-3-day old controls and mutants (5% and 14%, respectively; p=0.14, chi-square analysis). However, the incidence of sterility increases with age more in mutants than in controls, and by 4 weeks of age, significantly more mutant males (48%) fail to produce progeny than controls (27%; p=0.03). Overall, we found that no difference in fertility exists in 1-3-day- or 2-week-old flies, but that 4-week-old mutants are significantly less fertile than controls (Figure 4I), largely due to the pronounced increase in sterility in the population. This result confirms that *miR-317* is necessary for proper fertility in aging male flies.

### Relevant *miR-317* targets involved in mutant phenotypes

To investigate the molecular mechanism of *miR-317* involvement in GCTs, we used multiple online databases to compile a list of over 1000 predicted *miR-317* target genes (see Materials and Methods). 919 of these predicted targets were detected in our RNA-seq data, of which 142 are significantly dysregulated in *miR-317[ko]* testes (p<0.05; 88 downregulated and 54 upregulated; Supplemental Table 2). Although mostly highly significant, the levels of dysregulation of these genes are quite modest, with Log_2_ fold changes (Log_2_FC) ranging from 0.18-1.29 upregulated and -0.19 to -2.48 downregulated. This is consistent with previous results demonstrating that while each miRNA affects numerous targets, the change in expression of each mRNA target may be minor (Baek et al. 2008). Moreover, translational repression of a transcript by miRNAs can affect protein output without causing degradation of the mRNA; in this case, even effective miRNA targeting can yield little to no transcriptional changes.

Genes dysregulated in *miR-317[ko]* are expected to include both direct *miR-317* targets and indirectly affected, downstream genes. We placed the highest priority on upregulated genes because we expect transcripts that are directly targeted by *miR-317* to be more highly expressed in the mutant. Applying a stringent requirement that a gene should be upregulated with at least a Log_2_FC of 0.5 reduces the list of upregulated genes from 54 to 15 genes; we propose that these 15 genes are strong candidates to be direct targets of *miR-317* in testes (Table 1; Figure 2A).

**Table 1:** Predicted *miR-317* targets upregulated in testes in *miR-317[ko]* vs control; p<0.05 and Log_2_FC>0.5.

| Gene ID | Name | Protein Description | Log<sub>2</sub> Fold Change | Adjusted p value |
| --- | --- | --- | --- | --- |
| FBgn0037427 | *Osi17* | Osiris 17 ; integral component of plasma membrane | 1.29 | 4.12E-09 |
| FBgn0037419 | *Osi12* | Osiris 12 ; integral component of plasma membrane | 1.22 | 2.53E-08 |
| FBgn0039178 | *CG6356* | CG6356 ; transmembrane transport | 1.05 | 2.63E-11 |
| FBgn0066114 | *GlcAT-I* | GlcAT-I ; Golgi membrane | 1.02 | 9.05E-11 |
| FBgn0036661 | *CG9705* | Cold shock domain-containing protein CG9705 ; regulation of transcription, DNA-templated | 0.81 | 1.03E-21 |
| FBgn0037845 | *CG14694* | IP11787p ; transmembrane transporter activity | 0.79 | 1.62E-04 |
| FBgn0267337 | *rn* | Rotund ; transcription factor activity | 0.69 | 3.58E-11 |
| FBgn0031001 | *CG7884* | CG7884, isoform B ; no biological data available | 0.68 | 1.44E-03 |
| FBgn0034789 | *PIP5K59B* | PIP5K59B ; Phosphatidylinositol phosphate kinase activity | 0.67 | 3.30E-03 |
| FBgn0036725 | *CG18265* | GH10077p ; nucleic acid binding | 0.67 | 2.46E-03 |
| FBgn0037555 | *Ada2b* | Transcriptional Adaptor 2b ; regulation of transcription, DNA-templated | 0.61 | 6.22E-07 |
| FBgn0039064 | *CG4467* | CG4467, isoform B ; proteolysis | 0.58 | 2.93E-02 |
| FBgn0037837 | *CG14693* | Uncharacterized protein, isoform E | 0.56 | 3.38E-08 |
| FBgn0031275 | *GABA-B-R3* | Metabotropic GABA-B receptor subtype 3 ; G-protein coupled receptor signaling pathway | 0.54 | 1.98E-08 |
| FBgn0033476 | *oys* | Oysgedart ; lysophospholipid acyltransferase activity | 0.53 | 1.46E-09 |

We next sought to recapitulate *miR-317[ko]* mutant phenotypes via the upregulation of putative *miR-317* targets. For seven of the candidates in Table 1, we were able to acquire Drosophila stocks predicted to cause Gal4-induced OE (Table 2). For germline OE, we used *nanos-Gal4* (*nos-Gal4*), which is expressed in GSCs and early germ cells, and for somatic OE, we used *C587-Gal4*, which is expressed in cyst stem cells and cyst cells (Kawase et al. 2004) (Supplemental Figure 1, G and H). First, we overexpressed in germ cells those genes with UAS lines available and were unable to detect the AE phenotype or the formation of GCTs (Table 2). Next, we overexpressed in somatic cells using either *C587-Gal4* or *C587-Gal4; UAS-dCas9.VPR*, as appropriate (Table 2). The AE phenotype was not observed in any of the OE experiments. Notably, however, somatic expression of either *rotund* (*rn*) or *CG18265* is sufficient to cause GCTs (Table 2; Figure 5, A-C). Both *rn* and *CG18265* encode transcriptional regulators with no previously known roles in tumorigenesis. We conclude that OE of several *miR-317* candidate target genes can contribute to the formation of GCTs in the testis and propose that in wild-type testes, the miRNA *miR-317* functions as a tumor suppressor by repressing multiple targets..

**Figure 5.**
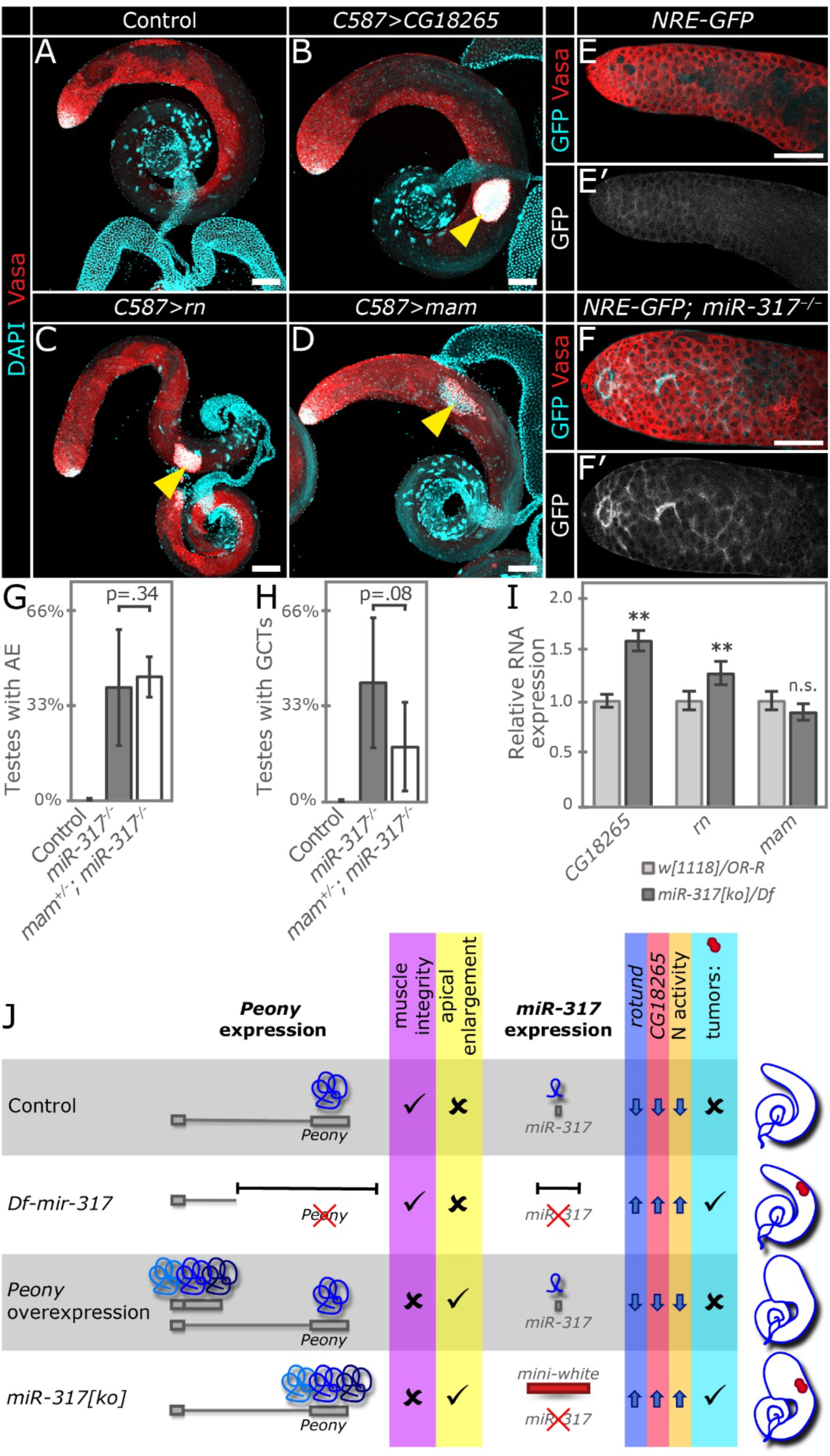
*miR-317* regulates multiple targets to suppress tumor formation. **A-D** Testes stained for DAPI (cyan) and Vasa (red). **A** Control testis. **B-D** Testes overexpressing *CG18265* (B), *rn* (C), or *mam* (D) under the control of the cyst-cell driver *C587-Gal4*, each exhibiting a GCT (yellow arrowheads). **E** Apex of control testis (*NRE-GFP*) in which GFP is expressed under the control of a Notch reporter element (NRE). GFP is nearly undetectable (shown in grayscale in E′). **F** GFP expression is strongly upregulated in *NRE-GFP; miR-317^-/-^* testes (shown in grayscale in F′). **G-H** Quantification of the percentage of testes that exhibit AE (G) or GCTs (H) in control, *miR-317^-/-^,* or *mam^+/-^; miR-317^-/-^* testes. Graphs depict mean ± standard deviation based on 3-7 biological replicates. **I** Results of qRT-PCR experiments assaying expression levels of *CG18265*, *rn*, and *mam* in male L3 larvae. Graph depicts mean ± confidence intervals based on 1 SEM. **J** Summary of our proposed model of the differential effects of *Peony* and *miR-317* on testes. In controls, *Peony* is expressed at its endogenous level, muscle integrity is intact, and AE is absent; *miR-317* is expressed, repressing target genes *rotund* and *CG18265* and Notch (N) activity, and tumors are absent. In *Df-miR-317* and *miR-317[ko]*, *miR-317* is absent, target genes are upregulated, N activity is increased, and GCTs (red clusters in testis schematics) form. In both *Peony* overexpression and *miR-317[ko], Peony* levels are high, causing loss of muscle integrity and the appearance of AE. **A-F**, scale bars 100 µm. **G-I**, statistical significance assessed using the Student’s t-test; **, p<.01.

**Table 2:**
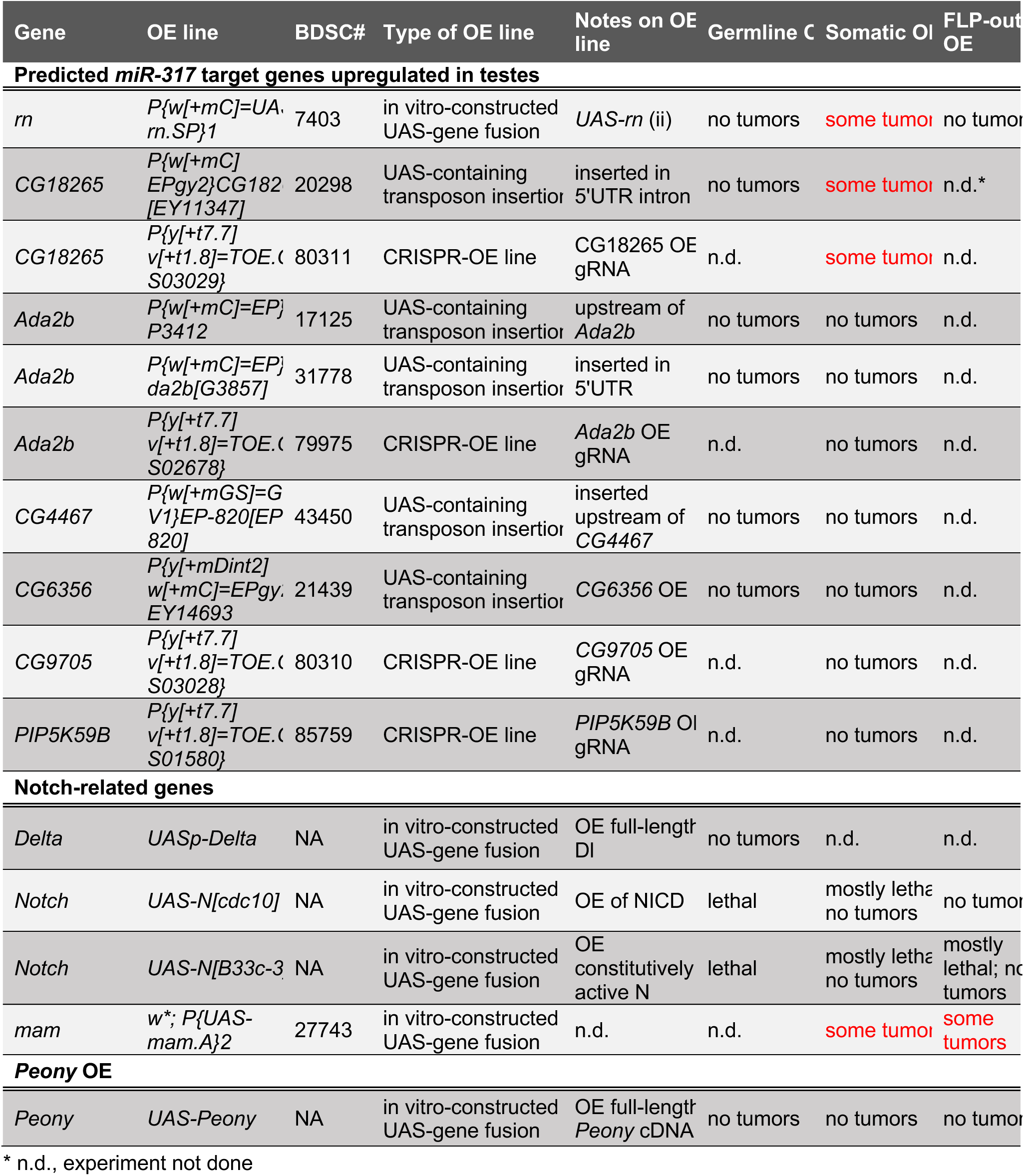
Testing for GCT formation caused by overexpression (OE) of candidate genes Predicted *miR-317* target genes upregulated in testes.

### The Notch cofactor Mastermind is implicated in the formation of GCTs

Among the genes predicted to be targeted by *miR-317* are many interactors and regulators of Notch signaling (Supplemental Table 3). To address the involvement of Notch in GCT formation, we attempted to augment Notch signaling strength by overexpressing Notch pathway components. We performed germline OE of the Notch ligand Delta or somatic OE of multiple forms of activated Notch. We also overexpressed Notch clonally using the Flp-out method (Struhl and Basler 1993). While Notch OE consistently causes a high degree of lethality, none of these conditions is sufficient to cause the formation of GCTs. We noted that *mastermind* (*mam*), which encodes a transcriptional cofactor in canonical Notch signaling, is among the transcripts with predicted *miR-317* binding sites (Supplemental Table 3), although it is not upregulated in our RNA-seq dataset (Supplemental Table 1). In another attempt to increase Notch signaling strength, we overexpressed *mam* in somatic cells using *C587-Gal4* and found that this condition results in formation of GCTs but not AE (Figure 5D). Furthermore, OE of *mam* in Flp-out clones results in tumors, similar to its somatic OE (Table 2).

To further investigate a potential involvement of the Notch pathway in *miR-317* LOF phenotypes, we tested Notch signaling strength using a reporter in which GFP is under the transcriptional control of a Notch responsive element (*NRE-GFP*) (Saj et al. 2010). We found that the GFP signal is nearly undetectable in most control testes from young, 1-day-old flies. In those that express detectable GFP, the pattern is consistent with weak Notch signaling activity in cyst cells (Figure 5, E and E′). Upon *miR-317* LOF, GFP expression is strongly upregulated in mature cyst cells while remaining undetectable in the cells closest to the apex (Figure 5, F and F′). We conclude that *miR-317* normally acts to repress Notch signaling in cyst cells.

Next, we tested whether Mam is necessary for either the GCT or the AE phenotype by introducing a strong *mam* LOF allele, *mam[8],* into the *miR-317[ko]* background. We found that the loss of one copy of *mam* fails to modify the AE phenotype (p=0.34; Figure 5G). In contrast, the incidence of GCTs is marginally suppressed (p=0.08) by *mam* heterozygosity (Figure 5H). Taken together, our results strongly suggest that Mam is both necessary and sufficient for testis GCT formation. Furthermore, as Mam is a crucial Notch cofactor, these results also raise the possibility that increased Notch signaling strength could serve to promote GCT formation in *miR-317* mutant testes.

Finally, to check the levels of *CG18265* and *rn*, the identified *miR-317* targets, during development, we isolated RNA from L3 larval males and subjected it to qRT-PCR. We also assayed the levels of *mam* in these animals. Consistent with our RNA-seq results (Supplemental Table 1), we found that *CG18265* and *rn* are significantly upregulated in *miR-317* mutant L3 larvae (Figure 5I). Also consistent with RNA-seq, we found that *mam* is not upregulated in mutant larvae, implying that despite its *miR-317* binding site, *mam* is likely not a direct target of the miRNA (Figure 5I).

Our data demonstrate that two noncoding RNAs encoded by the same genomic locus have distinct and separable functions: *miR-317* acts as a tumor suppressor by repressing multiple targets, whereas *Peony* regulates testis morphology. Upregulation of the *Peony* transcript causes lesions to appear in the testis muscle sheath leading to AE, and loss of *miR-317* causes upregulation of *CG18265* and *rn*, increased Notch signaling strength, and formation of GCTs (Figure 5J). Consistent with these conclusions, OE of *CG18265*, *rn*, or *mam* causes the formation of GCTs but does not lead to muscle disruption or AE. Ubiquitous OE of *Peony* from our inducible transgene does not cause GCT formation but results in a testis muscle breakage phenotype as well as a concomitant AE similar to that seen in *miR-317[ko]* testes (Figure 5J).

## Discussion

### Regulation of miRNA expression by differential polyadenylation

We demonstrate here that *Peony* and *miR-317* comprise a polycistronic unit and mutually affect one another’s expression levels. We identified a proximal polyA site near the annotated end of *Peony* and show evidence for another site, not yet molecularly defined, downstream of *miR-317*. We propose that transcriptional read-through of the proximal polyA site is required for expression of the miRNA, and the choice of polyA site determines whether *Peony* alone is expressed, or *miR-317* as well. Indeed, in the event that the polyA site is bypassed and *miR-317* is transcribed, pre-miRNA processing and cleavage would be predicted to de-stabilize the 5′ portion of the transcript that includes *Peony*. Therefore, expression of the two genes could in principle be mutually exclusive: proximal polyadenylation stabilizes the *Peony* transcript but prevents or reduces expression of *miR-317*, while transcriptional read-through allows miRNA processing but causes degradation of *Peony*.

It has been suggested that *miR-317* is expressed as a cluster with two downstream miRNA genes, *miR-277* and *miR-34*, both of which are processed from the lncRNA gene *CR43459* (Kadener et al. 2009; Zhou et al. 2018). Our data suggest that this is not the case, at least in testes. First, although our RNA-seq data demonstrate that *Peony* is strongly upregulated in *miR-317[ko]* testes, expression levels of *CR43459* do not change (Supplemental Table 1). Additionally, a TSS has been identified at the beginning of each of these ncRNA genes, *Peony* and *CR43459*, consistent with their independent transcription in spite of their close proximity (Zhou et al. 2018). Moreover, a genome-wide analysis of primary miRNAs in humans and mice suggests that a widespread mechanism leading to differential transcription of clustered miRNAs is through the use of alternative promoters (Chang et al. 2015).

In light of our finding that the functional lncRNA *Peony* is adjacent to and co-transcribed with *miR-317*, phenotypes that would ordinarily have been ascribed to the miRNA should be scrutinized carefully for involvement of *Peony*. For instance, several studies reported phenotypes that they attributed to loss of *miR-317*, when the allele used was a P{GT1} transposon (BG01900) inserted in *Peony*, 1730 bp 5′ of *miR-317* (Yamamoto et al. 2008; Edwards et al. 2009; Fricke et al. 2014). Moreover, care should be given in interpretation of phenotypes caused by the *miR-317[ko]* allele since, as demonstrated here, this allele results in a strong upregulation of *Peony*, which could produce confounding results. Potentially helpful in this respect is the new *Df-miR-317* allele generated for this work, which removes both *miR-317* and *Peony*, facilitating the unambiguous attribution of phenotypes to the miRNA.

An important and enticing question is whether other miRNA genes are regulated in a way similar to the *Peony-miR-317* locus. We carefully examined the genomic loci of all known Drosophila miRNA genes, of which there are over 260 (Supplemental Table 4; Supplemental Figure 4A). The most common genomic context for miRNA genes is presence in the intron of a coding gene (118/264 miRNA genes, 44.7%). Many others are located in exons or in intergenic space, while an appreciable minority (46/264 genes, 17.4%) reside closely upstream or downstream of adjacent genes in a way similar to the close proximity of *Peony* and *miR-317* (Supplemental Figure 4B). This 17.4% of miRNA genes, which we term “putatively co-transcribed,” represents an intriguing avenue for further investigation to determine if they are indeed co-expressed with adjacent genes and whether alternative polyadenylation may play a role in their differential expression.

### The cryptic yet crucial functions of lncRNAs

It has recently become appreciated that eukaryotic genomes undergo pervasive transcription, in stark contrast to the historical view that the transcriptome consists mainly of protein-coding transcripts and other RNAs involved primarily in translation: tRNAs, rRNAs, and a few others. This widespread transcription results in a vast number of non-coding transcripts, the functions of most of which are still unknown (Jensen et al. 2013). MiRNAs comprise one class of non-coding RNAs (ncRNAs), but while the canonical function of miRNAs is well-known, the same cannot be said for the lncRNAs. Indeed, although many thousands of these transcripts exist in animal cells, their functions remain mostly obscure. It is a subject of much debate whether many lncRNAs are the result of transcriptional noise with no bona fide functions within the cell. Counter to this proposition has been the elucidation of important functions for an ever-increasing number of lncRNA genes as well as clear instances of evolutionary conservation, which would be unexpected for transcripts with no importance (Guttman et al. 2009; Chen 2016; Li and Liu 2019). Indeed, the study of lncRNAs’ molecular and cellular functions and the determination of their roles in living cells remains an important and tantalizing problem in modern biology. Our finding that *Peony* is co-transcribed with an important tumor-suppressive miRNA and that the lncRNA itself also affects muscle integrity and testis morphology adds an important element to the growing body of evidence demonstrating the important roles of lncRNAs in numerous contexts.

### *MiR-317* regulates multiple genes to repress tumor formation in developing testes

MiRNAs or miRNA clusters can target multiple genes in the same pathway to ensure a speedy and robust response (Mestdagh et al. 2010; Uhlmann et al. 2012; Cicek et al. 2016; Hart et al. 2019). We report here that repression of at least two putative *miR-317* targets, *rn* and *CG18265*, is necessary to prevent the onset of tumors in Drosophila testes. The upregulation of both genes in *miR-317* mutants is modest (1.26 to 1.61-fold) as measured both by RNA-seq (Supplemental Table 1) and by qRT-PCR (Figure 5I). This is illustrative of the miRNA’s typical regulation of multiple targets, in which the effect is not a binary on/off result but rather a fine-tuning of gene expression (Baek et al. 2008), by which the miRNA can serve to reinforce cell fates, regulate responses to environmental cues, or buffer stochastic fluctuations in gene expression or signaling. We also note that the penetrance of the GCT formation phenotype is low and variable, in that most germline cysts escape tumorigenesis and apparently undergo spermatogenesis properly. In fact, in most experiments, the majority of males examined had no tumors. We propose that this reflects the effect of *miR-317*’s expression tuning on the remarkably robust processes of spermatogenesis, and that *miR-317* deficiency sensitizes the system such that these processes are marginally more prone to failure, leading to an infrequent tumorigenic event.

Both *rn* and *CG18265* encode C2H2 zinc-finger transcription factors. Interestingly, Rn is reported to affect Notch signaling in multiple contexts: *rn* mutation dominantly enhances the *split* phenotype caused by a recessive *Notch* allele (Brand and Campos-Ortega 1990); and the expression patterns of the Notch ligands Delta and Serrate are perturbed by *rn* mutation in imaginal discs (St Pierre et al. 2002). To our knowledge, no link between *CG18265* and Notch signaling has yet been demonstrated.

Rn has important roles in olfactory neuron specification and imaginal disc development. While not an essential gene, both male and female *rn* mutants are sterile. This sterility is not germline-dependent (Kerridge and Thomas-Cavallin 1988), suggesting that Rn has a role in the somatic gonadal cells. While it is not known why *rn* mutants are sterile, our findings suggest that *miR-317* acts in somatic cells of testes to modulate the levels or expression pattern of *rn* and prevent germline overproliferation.

Our data show that GCTs in *miR-317* LOF testes arise during development, such that they already exist in newly eclosed males, and our evidence suggests that new tumors do not form during adulthood. This is consistent with our finding that the *miR-317* targets *CG18265* and *rn* are upregulated in *miR-317* LOF L3 larvae (Figure 5I) and young adult testes (Supplemental Table 1), suggesting that the crucial time for *miR-317*’s repressive action in terms of tumor formation is during development. As metamorphosis is analogous to human puberty (Guirado et al. 2023), the timing of GCTs in *miR-317* LOF animals is reminiscent of the incidence of human seminoma, which is a common type of GCT in males that frequently manifests beginning in adolescence (Frazier et al. 2012). Seminomas are tumors composed of germ cells that remain in their undifferentiated state (Penn et al. 2018), contrasting with other testicular neoplasms such as teratomas, in which pluripotent cells are capable of giving rise to progeny of all three germ layers (Nettersheim et al. 2016). The *miR-317* mutant GCTs, similar to seminomas, appear to contain only undifferentiated germ cells.

### Sufficiency of Mam and Notch in driving GCT formation

We show in the present study that OE of Mam in cyst cells is sufficient to cause GCT formation, and reduction of Mam partially abrogates the incidence of GCTs. As Mam has been well-studied as an important transcriptional cofactor in canonical Notch signaling, these results suggest the intriguing possibility that increased Notch signaling strength is causative in GCT formation. However, it has also been reported that in various contexts, Mam functions in a Notch-independent manner (Kankel et al. 2007; Vied and Kalderon 2009; Zhang et al. 2011; Lobo-Pecellin et al. 2019). Our results do not allow us to distinguish between a canonical or Notch-independent function for Mam in tumor formation; therefore, a Notch-independent Mam function remains a possibility in the context of testis GCT formation. However, in light of our finding that *miR-317[ko]* causes increased Notch activity in cyst cells, the same cell type in which Mam OE results in GCTs, we favor a model in which cyst cell-expressed Mam participates in this Notch signaling.

While we were unable to show that overexpressed Notch drives GCT formation, the possibility remains that increased Notch signaling may drive tumorigenesis in conditions that we did not test. Further manipulation of the timing and strength of the Notch pathway would be required to truly exclude this possibility; additionally, a more detailed exploration of the genetic interaction between *Notch* and *miR-317* should prove informative. Alternatively, Notch OE may not induce GCTs if Mam is a limiting factor in determining the strength of Notch signaling. In this instance, reducing the gene copy number of *mam* decreases the incidence of tumors as we demonstrate here, but increasing Notch without simultaneously increasing Mam is insufficient to induce tumor formation. Consistent with this possibility, in Drosophila ovaries, somatic OE of activated Notch and Mam simultaneously results in tumor-like overgrowth, while expression of Notch alone does not (Defreitas et al. 2020).

Intriguingly, the consequences of Notch involvement in various processes differ in a cell- and context-dependent manner. For instance, Notch can behave either as an oncogene or as a tumor suppressor, depending on the cell type; similarly, either proliferation or terminal differentiation can be promoted by increased Notch signaling (Koch et al. 2013; Yatsenko and Shcherbata 2018; Yatsenko and Shcherbata 2021; Zamfirescu et al. 2022). In Drosophila, OE of Notch in neural stem cells induces tumorigenesis, while in intestinal stem cell lineages, increased Notch is required for differentiation (Ohlstein and Spradling 2006; Wang et al. 2006; Ohlstein and Spradling 2007; Bowman et al. 2008).

In the Drosophila testis, Notch signaling was shown to be required for germ cell survival: Delta produced in the germline and Notch in the somatic cyst cells are both required to prevent cell death of the germ cells (Ng et al. 2019). In light of our results showing that increased Notch signaling in *miR-317* mutants coincides with tumor formation, it is possible that the levels of Notch components or the strength of Notch signaling must be maintained within a particular range to prevent either cell death or overproliferation. Another possibility is that cysts at different developmental stages have variable competence to undergo Notch-mediated germline overproliferation. It has been reported that Notch signaling is not active in the youngest cyst cells near the apex, rather only in more differentiated, slightly more basal cells (Ng et al. 2019). Thus, increased Notch signaling strength in earlier cyst cells in the spermatogonial region may have the propensity non-autonomously to cause overproliferation of their associated germ cells. Future studies will be required to assess the competence of different stages of cysts to undergo tumorigenesis.

## Declaration of interests

The authors declare that they have no competing interests. This study includes no data deposited in external repositories.

## Data Availability Statement

RNA-seq data will be deposited in the NCBI Gene Expression Omnibus (GEO) and made publicly available upon publication. Confocal microscopy images and quantitative image analyses will be provided by the corresponding author upon reasonable request. All *Drosophila melanogaster* mutant and transgenic lines generated in this study will be made available upon request and deposited to the Bloomington Drosophila Stock Centre following publication.

## Acknowledgments

We would like to thank the Bloomington Drosophila Stock Center (NIH P40OD018537) and the Kyoto Drosophila Stock Center at the Kyoto Institute of Technology for Drosophila stocks used in this study. We also thank the Developmental Studies Hybridoma Bank, created by the NICHD of the NIH and maintained at The University of Iowa, for antibodies. We are grateful to Dorothea Godt, Gerd Vorbrüggen, Hannele Ruohola-Baker, Herbert Jäckle, Renate Renkawitz-Pohl, Stefan Jakobs, and Christian Bökel for kindly sharing flies and reagents with us. We thank all Shcherbata lab members for critical reading of the manuscript and helpful suggestions; Rucha Hebalkar for assistance with expression pattern stains; and Melina Schwier and Robin Dewender for assistance with dissections and qRT-PCR. We are grateful to Dr. Frederic Bonnet and the MDI Biological Laboratory Light Microscopy Facility (RRID:SCR_019166). This research was supported by the German Research Foundation (DFG) grant numbers 521749003 and INST 192/615-1 and an Institutional Development Award (IDeA) from the NIGMS of the NIH under grant numbers P20GM103423 and P30GM154610.

## Author contributions

Conceptualization, TDC, HRS; Methodology, TDC, HRS; Investigation, TDC; Writing, TDC, HRS; Visualization, TDC, HRS; Funding Acquisition, HRS.

## Materials and Methods

### Fly Stocks and Genetics

Flies were maintained at 25°C at constant humidity and a 12h-12h light-dark cycle. They were kept on standard cornmeal-yeast-agar fly food. All genotypes used are listed in Key Resources Table. For experiments utilizing *Tub-Gal80[ts],* crosses were performed and progeny were allowed to develop for 3 days at 25°C, at which temperature Gal4 activity is low in the presence of Gal80[ts]. After 3 days, larvae were shifted to 29°C to allow Gal4-induced expression until dissection (L3 or adult).

The following stocks were provided by the Bloomington Drosophila Stock Center (BDSC): *w[1118]* (BDSC 6326); *OR-R* (BDSC 5); *C587-Gal4* (BDSC 67747); *P{UAS-3xFLAG.dCas9.VPR}attP40* (BDSC 66561); *P{w[+mC]=EP} EP3412* (BDSC 17125); *P{w[+mC]=EP}Ada2b[G3857]* (BDSC 31778); *w[*]; P{w[+mW.hs]=GawB}bab1[Pgal4-2]/TM6B, Tb[1]* (BDSC 6803); *P{w[+mC] EPgy2}CG18265[EY11347]* (BDSC 20298); *P{y[+t7.7] v[+t1.8]=TOE.GS02678}* (BDSC 79975); *P{y[+t7.7] v[+t1.8]=TOE.GS03029}* (BDSC 80311); *P{w[+mGS]=GSV1}EP-820[EP-820]* (BDSC 43450); *P{y[+t7.7] v[+t1.8]=TOE.GS03028}* (BDSC 80310); *Mi{MIC}MI02131* (BDSC 37146); *P{w[+mW.hs]=GawB}elav[C155]* (BDSC 458); *P{y[+mDint2] w[+mC]=EPgy2}EY14693* (BDSC 21439); *w*; P{UAS-mam.A}2* (BDSC 27743); *mam[8]/CyO* (BDSC 1596); *miR-317[ko]* (BDSC 58926); *P{w[+mW.hs]=GawB}MJ12a* (BDSC 6991); *w[1118]; P{w[+m*]=NRE-EGFP.S}5A* (BDSC 30727); *P{y[+t7.7] v[+t1.8]=TOE.GS01580}* (BDSC 85759); *P{w[+mC]=UAS-rn.SP}1* (BDSC 7403); *RpII215[c4]* (BDSC 3663); *y1 w*; P{tubP-GAL4}LL7/TM3, Sb1 Ser1* (BDSC 5138); *w[*]; P{w[+mC]=tubP-GAL80[ts]}10* (BDSC 7108); and *w[*]; P{w[+mC]=tubP-GAL80[ts]}2* (BDSC 7017). The following stocks were provided by the Kyoto Drosophila Stock Center at the Department of Drosophila Genomics and Genetic Resources (DGGR), Kyoto Institute of Technology: *P{GSV6}GS11364* (DGGR 203275); *y* w*; P{w+mW.hs=GawB}NP1624* (DGGR 104055); and *P{w+mC=vas-GAL4.2.6}2* (DGGR 109996). The following stocks were procured from the Exelixis collection at Harvard Medical School: *PBac{WH}f04665* and *P{XP}d02752*.

Among the stocks used for OE of putative *miR-317* targets were CRISPR-OE lines. These stocks ubiquitously expresses a guide RNA (gRNA) specific for a gene of interest; when crossed to a stock containing both an appropriate Gal4 driver and *UAS-dCas9.VPR*, a construct expressing a nuclease-dead Cas9 protein fused with a transcriptional activation domain, this can cause the OE of the gene of interest from its endogenous locus (Lin et al. 2015). Other stocks used were: *UAS-Peony* (this study); *Df-miR-317* (this study); *P{w+mC=bam-GFP.-799/+133}3LR* (Chen and McKearin 2003); *P[UASp-Delta-2]* (Ward et al. 2006); *w*; P{GAL4-nos.NGT}40; P{GAL4::VP16-nos.UTR}CG6325MVD1* (a gift from Hannele Ruohola-Baker); *Mef2-Gal4* (a gift from Gerd Vorbrüggen); *Tub-Gal4* (chromosome 2; a gift from Stefan Jakobs); and *zfh1::T2A::Gal4* (a gift from Christian Bökel). Alleles used to express activated forms of Notch were *UAS-N[cdc10]*, which expresses an intracellular fragment of Notch (Brennan et al. 1999) and *UAS-N[B33c-3]*, which results in the expression of a constitutively active form of Notch lacking most of the extracellular domain (Larkin et al. 1996).

### Fertility assay

Fertility assay was performed as previously reported (Herrera et al. 2021). Individual control (*w[1118]/OR-R*) or mutant (*miR-317[ko]/Df*) males of either 0-3 days, 2 weeks, or 4 weeks of age were crossed to two 1-3-day-old control virgin females. For each time point, 50 males were initially crossed. Flies were allowed to mate for 48 hours, then they were removed from the vials. Any crosses in which the male or either female died within the 48-hour period were not included in the analysis. After 10-12 days, the progeny from each cross were counted as a measure of each male’s fertility.

### Immunofluorescence and confocal microscopy

Adult flies were dissected with Dumont #5 forceps (Fine Science Tools) in cold phosphate-buffered saline (PBS). Testes were fixed in 4% formaldehyde (Polysciences 18814-20) in PBT (PBS + 0.2% Triton-X 100 (Sigma 108603)) for 20 minutes, then rinsed in PBT multiple times for a total of at least 30 minutes. Then, testes were blocked in PBTB solution (PBT + 5% normal goat serum (Abcam ab7481) and 0.1% bovine serum albumin (Applichem A1391)) for at least 30 minutes. Next, testes were incubated in a solution of primary antibodies diluted in PBTB overnight at 4°C. After primary antibody incubation, testes were rinsed multiple times for at least 30 minutes, blocked in PBTB for at least 30 minutes, and incubated in a solution of secondary antibodies diluted in PBTB, either at room temperature for 2.5 hours or at 4°C overnight. Then, testes were rinsed multiple times in PBT, incubated in a 1 μg/mL solution of DAPI in PBT for 10 minutes, and rinsed three more times with PBT. Finally, the buffer was removed and testes were stored in Vectashield (Vector Labs) until being mounted on slides and imaged.

The following mouse monoclonal antibodies were procured from the Developmental Studies Hybridoma Bank and used at the dilutions shown: anti-Adducin (1:20), anti-LaminC (1:20), anti-Dlg (1:20), anti-Eyes absent (1:100), anti-Fas3 (1:20), and anti-β-Galactosidase (1:200). Other antibodies used were: chicken anti-GFP (1:5000, Abcam); rabbit anti-mCherry (1:200, Abcam); rabbit anti-Phosphohistone H3 (1:10000, Upstate Biotechnology); guinea pig anti-Traffic jam (1:5000, a gift from Dorothea Godt); guinea pig anti-β3-Tubulin (1:1500, a gift from Renate Renkawitz-Pohl); and rabbit anti-Vasa (1:5000, a gift from Herbert Jäckle). Tissues were also counterstained using the DNA stain DAPI (VWR) to mark nuclei and Cy3-labeled phalloidin (ThermoFisher) to mark actin filaments. Fluor-conjugated secondary antibodies were obtained from Jackson Immunoresearch or ThermoFisher and used at a 1:500 dilution.

Imaging at Hannover Medical School was performed using a Zeiss LSM 700 confocal microscope equipped with dual photomultiplier tubes (PMTs); four laser lines at 405, 488, 555, and 647 nm; and the following objectives: 10x air (numerical aperture (NA) 0.45), 25x oil (NA 0.8), and 40x oil (NA 1.3). Imaging at the Mount Desert Island Biological Laboratory was performed using a spinning-disk confocal (CSU-W1, Yokogawa, Japan) on a Nikon inverted Ti-Eclipse microscope stand (Nikon Instruments Inc., Japan), using a Nikon 10x air objective (NA 0.45; ref: MRD70170). DAPI and AF488 fluorescence were excited using a 405 nm and 488 nm lasers and collected using bandpass filters (Chroma ref: ET436/20x and Semrock ref: FF01-525/50). Images were acquired in 1024x1024 pixels with a Scientific CMOS Zyla 4.2 (Andor Technology, United Kingdom) controlled with NIS AR 5.41.02 software (build 1711, Nikon Instruments Inc., Japan) and saved in nd2 file format. Z-stack images were taken of all samples. Either single focal plane or z-projections of multiple focal planes were used in figures, as appropriate to best depict each phenotype. Images were processed using Fiji software (Schindelin et al. 2012) and assembled in Adobe Illustrator (Adobe, Inc.).

### miRNA target prediction

The following databases were used to search for predicted targets of *miR-317*: TargetScanFly (http://www.targetscan.org), MinoTar (https://www.flyrnai.org/cgi-bin/DRSC_MinoTar.pl), and microRNA.org (http://www.microrna.org). 975 targets were predicted by microRNA.org, 31 by MinoTar, and 102 by TargetScanFly; 76 targets were predicted by multiple databases for a total of 1030 predicted targets.

### qRT-PCR

Drosophila tissues were dissected in cold PBS and transferred to microcentrifuge tubes. Total RNA was prepared using a standard Trizol-chloroform extraction followed by an isopropanol precipitation. RNA was quantified using a Nano-drop spectrophotometer (ThermoFisher) and treated with DNaseI. For expression levels of mRNA, lncRNA, or pre-miRNA, SYBR Green-based qPCR was performed. For measuring mature miRNA levels, a TaqMan-based assay was performed. Reactions were run in triplicate. Either α-Tubulin 84B (SYBR Green qPCR) or 2S rRNA (TaqMan qPCR) was used as an endogenous control. Expression fold change (FC) was determined using the expression FC = 2^-ΔΔCt^.

### 3′ RACE PCR

To identify the site of polyadenylation between *Peony* and *miR-317*, a 3′ RACE procedure was performed. Total RNA was extracted from five adult male flies of the genotype *RpII215[c4]* using TRIzol-chloroform extraction and isopropanol precipitation. Briefly, the flies were homogenized in 400 µL of TRIzol, 80 µL of chloroform was added, and the sample was shaken vigorously by hand. The sample was centrifuged for 15 minutes at 4°C and 12,000 x g. The upper, aqueous phase was moved to a new tube, and isopropanol was used to precipitate RNA at room temperature for 10 minutes. The sample was centrifuged at 4°C and 12,000 x g for 10 minutes to pellet RNA. The pellet was washed with 75% ethanol, dried, and suspended in RNAse-free water. 4 µg of RNA was used for each reverse transcription reaction.

Reverse transcription was performed using SuperScript II (Invitrogen) according to the manufacturer’s instructions. A control reaction lacking reverse transcriptase (-RT) was performed in parallel with the experimental reaction. An anchored oligo(dT) primer appended with a 5′ adapter sequence (5′-GGCCACGCGTCGACTAGTAC(T_17_)V-3′; Integrated DNA Technologies, IDT) was used for the RT reaction. The RT reaction progressed for two hours at 42°C, followed by 15 minutes at 95°C to inactivate the polymerase.

Identification of the polyA site was performed using a semi-nested PCR protocol. First, an “outer” nested reaction was performed using a gene-specific primer (*RACE_F1*, 5′-TTCGTTCGCTTGTGAGTGTCC-3′) within the second exon of *Peony*, and as reverse primer, the adapter portion of the anchored oligo(dT) was used: *3’ RACE PCR* (5′-GGCCACGCGTCGACTAGTAC-3′; IDT). This reaction proceeded for 25 cycles of PCR. Next, this PCR product was diluted 1:100 in water and used as template for the “inner” nested PCR. For this reaction, a forward primer (*RACE_F2*, 5′-ACACATACACGCACACAAACGG-3′) was used that is downstream of *RACE_F1*, still within the second exon of *Peony*. The same reverse primer, *3’ RACE PCR* (IDT) was used. PCR was performed for 35 cycles in three 50 µL reactions. The PCR product was run on a 1% agarose gel, and a prominent band at 400 bp was excised from the gel and the DNA isolated using a gel-extraction kit (Zymo). This DNA was Sanger-sequenced using the *RACE_F2* primer.

### Generation of *UAS-Peony*

The pUASt-attB vector was linearized with XhoI. The *Peony* cDNA was generated from RNA taken from *w[1118]* flies and PCR amplified using the following primers (lowercase bases indicate overlap to the vector; uppercase bases correspond to genomic sequence): *UAS-Peony_F* aacagatctgcggccgcggcGAATTCTGAACACAGCAAAAAGTACTGCATATAATAC; *UAS-Peony_R2* aagatcctctagaggtacccGCTCAGCAGAGACACAAACCG. Gibson assembly was used to assemble insert and vector. Transgenesis was performed by GenetiVision (Houston, Texas); the transgene was site-specifically integrated at the VK1 attP site on chromosome 2R.

### RNA-sequencing

RNA was extracted from testes from young, 0-1-day-old *w[1118]* and *miR-317[ko]* flies. Accessory glands and other components of the male reproductive system were separated from testes prior to RNA extraction. All samples were in triplicate. Total RNA was treated with DNaseI and submitted to the Transcriptome and Genome Analysis Laboratory (TAL) at the Georg-August-University, Göttingen, Germany. RNA quality was assessed by TAL prior to library preparation and sequencing. Sequencing was performed on an Illumina HiSeq4000 system, generating 50-bp reads and approximately 50 million reads per sample. Alignment and differential expression analysis was performed by TAL, resulting in the identification of 1844 genes in testes that were dysregulated in *miR-317[ko]* (Supplemental Table 1).

### Reagent Table

| Reagent type | Source | Identifier | Gene |
| --- | --- | --- | --- |
| Drosophila stock | BDSC #5 | *OR-R* | - |
| Drosophila stock | BDSC #67747 | *C587-Gal4* | - |
| Drosophila stock | BDSC #66561 | *P{UAS-3xFLAG.dCas9.VPR}attP40* | - |
| Drosophila stock | BDSC #17125 | *P{w[+mC]=EP}EP3412* | *Ada2b* |
| Drosophila stock | BDSC #31778 | *P{w[+mC]=EP}Ada2b[G3857]* | *Ada2b* |
| Drosophila stock | BDSC #79975 | *P{y[+t7.7] v[+t1.8]=TOE.GS02678}* | *Ada2b* |
| Drosophila stock | BDSC #6803 | *w[\*]; P{w[+mW.hs]=GawB}bab1[Pgal4-2]/TM6B, Tb[1]* | *bab1-Gal4* |
| Drosophila stock | BDSC #20298 | *P{w[+mC] EPgy2}CG18265[EY11347]* | *CG18265* |
| Drosophila stock | BDSC #80311 | *P{y[+t7.7] v[+t1.8]=TOE.GS03029}* | *CG18265* |
| Drosophila stock | BDSC #43450 | *P{w[+mGS]=GSV1}EP-820[EP-820]* | *CG4467* |
| Drosophila stock | BDSC #21439 | *P{y[+mDint2] w[+mC]=EPgy2}EY14693* | *CG6356* |
| Drosophila stock | BDSC #80310 | *P{y[+t7.7] v[+t1.8]=TOE.GS03028}* | *CG9705* |
| Drosophila stock | BDSC #37146 | *Mi{MIC}MI02131* | *Peony* |
| Drosophila stock | BDSC #458 | *P{w[+mW.hs]=GawB}elav[C155]* | *elav-Gal4* |
| Drosophila stock | BDSC #27743 | *w\*; P{UAS-mam.A}2* | *mastermind* |
| Drosophila stock | BDSC #1596 | *mam[8]/CyO* | *mastermind* |
| Drosophila stock | BDSC #58926 | *miR-317[ko]* | *miR-317* |
| Drosophila stock | BDSC #6991 | *P{w[+mW.hs]=GawB}MJ12a* | *MJ12a-Gal4* |
| Drosophila stock | BDSC #30727 | *w[1118]; P{w[+m\*]=NRE-EGFP.S}5A* | *NRE-GFP* |
| Drosophila stock | BDSC #85759 | *P{y[+t7.7] v[+t1.8]=TOE.GS01580}* | *PIP5K59B* |
| Drosophila stock | BDSC #7403 | *P{w[+mC]=UAS-rn.SP}1* | *rotund* |
| Drosophila stock | BDSC #3663 | *RplI215[c4]* | *RplI215* |
| **Drosophila stock** | BDSC #5138 | *y1 w\**; *P{tubP-GAL4}LL7/TM3, Sb1 Ser* | *Tub-Gal4 (III)* |
| **Drosophila stock** | BDSC #7108 | *w[\*]; P{w[+mC]=tubP-GAL80[TS]}10* | *Tub-Gal80[ts] (I)* |
| **Drosophila stock** | BDSC #7017 | *w[\*]; P{w[+mC]=tubP-GAL80[TS]}2* | *Tub-Gal80[ts] (I)* |
| **Drosophila stock** | BDSC #6326 | *w[1118]* | *white* |
| **Drosophila stock** | DGGR #203275 | *P{GSV6}GS11364* | *Peony* |
| **Drosophila stock** | DGGR #104055 | *y\* w\**; *P{w+mW.hs=GawB}NP1624* | *tj-Gal4* |
| **Drosophila stock** | DGGR #109996 | *P{w<sup>+mC</sup>=vas-GAL4.2.6}2* | *vasa-Gal4* |
| **Drosophila stock** | Exelixis #f04665 | *PBac{WH}f04665* | *Peony* |
| **Drosophila stock** | Exelixis #d02752 | *P{XP}d02752* | *Peony* |
| **Drosophila stock** | this study | *UAS-Peony* | *Peony* |
| **Drosophila stock** | this study | *Df-miR-317* | *miR-317* |
| **Drosophila stock** | (Brennan et al, 1999) | *UAS-N[cdc10]* | *Notch* |
| **Drosophila stock** | (Chen & McKearin, 2003) | *P{w+mC=bam-GFP.-799/+133}3LR* | *bam* |
| **Drosophila stock** | (Larkin et al, 1996) | *UAS-N[B33c-3]* | *Notch* |
| **Drosophila stock** | (Ward, et al. 2006) | *P[UASp-Delta-2]* | *Delta* |
| **Drosophila stock** | Gift from Hannele Ruohola-Baker | *w\**; *P{GAL4-nos.NGT}40; P{GAL4::VP1, nos.UTR}CG6325MVD1* | *nos-Gal4* |
| **Drosophila stock** | Gift from Gerd Vorbrüggen | *Mef2-Gal4* | *Mef2-Gal4* |
| **Drosophila stock** | Gift from Stefan Jakob | *Tub-Gal4* | *Tub-Gal4 (II)* |
| **Drosophila stock** | Gift from Christian Bökel | *zfh1:T2A:Gal4* | *zfh1* |
| **Antibody** | DSHB | Mouse anti-Adducin 1:20 | NA |
| **Antibody** | DSHB | Mouse anti-Eyes absent 1:100 | NA |
| **Antibody** | DSHB | Mouse anti-β-Galactosidase 1:200 | NA |
| **Antibody** | DSHB | Mouse anti-LaminC 1:20 | NA |
| **Antibody** | Abcam | Chicken anti-GFP 1:5000 | NA |
| **Antibody** | Abcam | Rabbit anti-mCherry 1:200 | NA |
| **Antibody** | Upstate Biotechnology | Rabbit anti-Phosphohistone H3 1:10000 | NA |
| **Antibody** | Gift from Dorothea Godt | Guinea pig anti-Traffic jam 1:5000 | NA |
| **Antibody** | Gift from Renate Renkawitz-Pohl | Guinea pig anti-β3-Tubulin 1:1500 | NA |
| **Antibody** | Gift from Herbert Jäck | Rabbit anti-Vasa 1:5000 | NA |
| **Stain** | VWR | DAPI | NA |
| **Stain** | Thermofisher | Cy3-labeled Phalloidin | NA |
| **DGGR = Department of Drosophila Genomics and Genetic Resources, Kyoto Institute of Technology** |  |  |  |
| **BDSC = Bloomington Drosophila Stock Center** |  |  |  |
| **DSHB = Developmental Studies Hybridoma Bank** |  |  |  |

## Supplementary Information

**Supplemental Figure 1.**
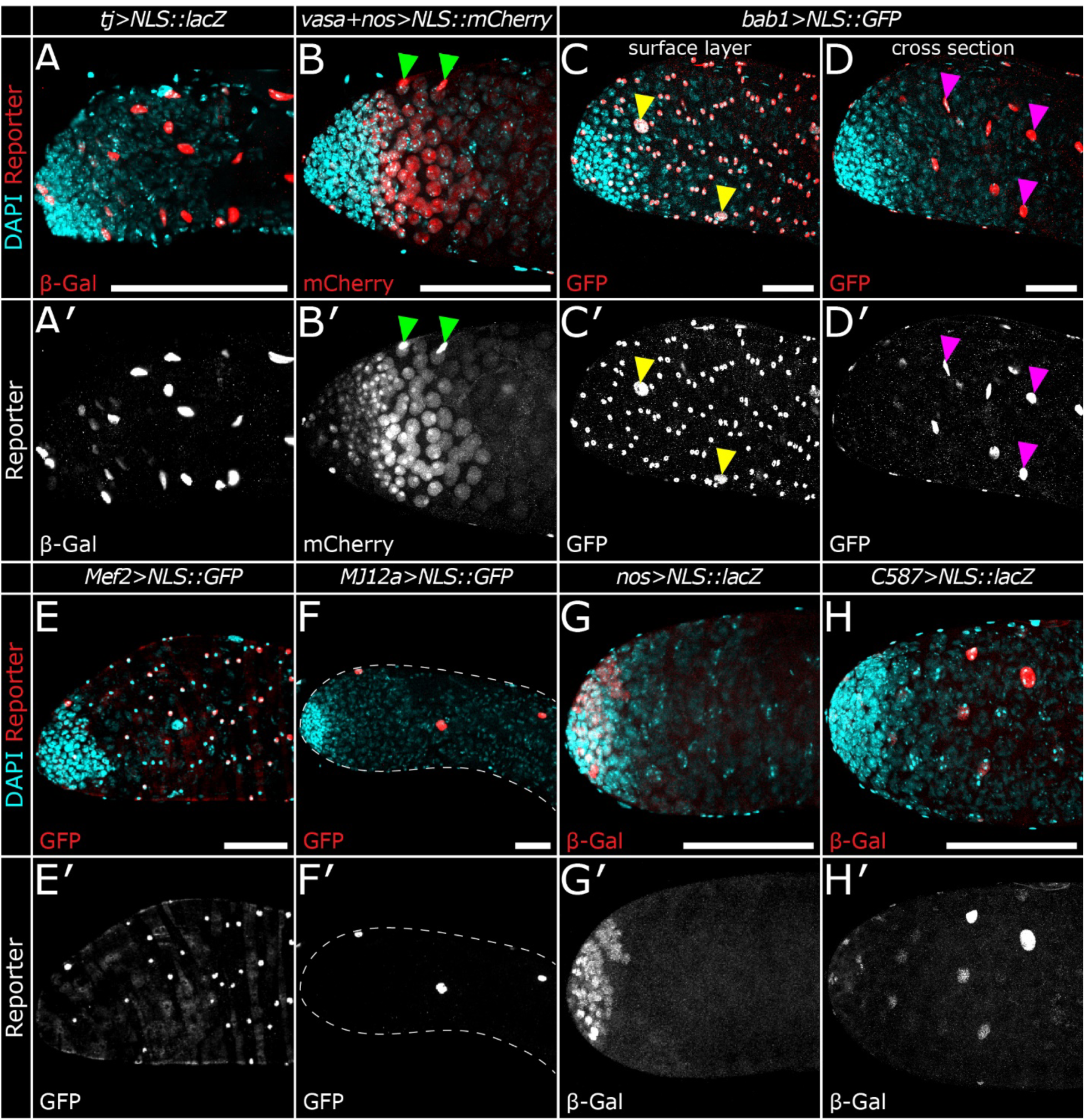
Testis expression patterns of Gal4 lines used in this study. **A-D** Expression patterns of Gal4 lines visualized by expression of reporters from UAS transgenes bearing a nuclear localization sequence (NLS): *UAS-NLS::lacZ* (A, G, & H); *UAS-NLS::mCherry* (B); and *UAS-NLS::GFP* (C-F). A′-H′ show reporter protein expression in single grayscale panels. **A** *traffic jam-Gal4* (*tj-Gal4*) is expressed in somatic cyst cells. **B** *vasa-Gal4; nos-Gal4* (*vasa+nos-Gal4*) is expressed primarily in germline cells but exhibits expression in some somatic cyst cells as well (examples: green arrowheads). **C-D** *bric a brac 1-Gal4* (*bab1-Gal4*) is expressed in somatic cells of the testis. Panel C shows a surface section in which muscle cell nuclei (numerous small nuclei) and pigment cell nuclei (yellow arrowheads) express GFP. D shows a deeper, internal cross section in which somatic cyst cell nuclei express GFP (examples: magenta arrowheads). **E** *Mef2-Gal4* drives expression in some, but not all, muscle cell nuclei. **F** *MJ12a-Gal4* is expressed only in pigment cells. **G** *nanos-Gal4* (*nos-Gal4*) is expressed in early germline cells near the apex. **H** *C587-Gal4* is expressed in somatic cyst cells. A, C, D, and F each depicts a single focal plane. Panels B, E, G, and H each depicts a z-projection of multiple focal planes to better illustrate expression patterns. All scale bars: 100 µm.

**Supplemental Figure 2.**
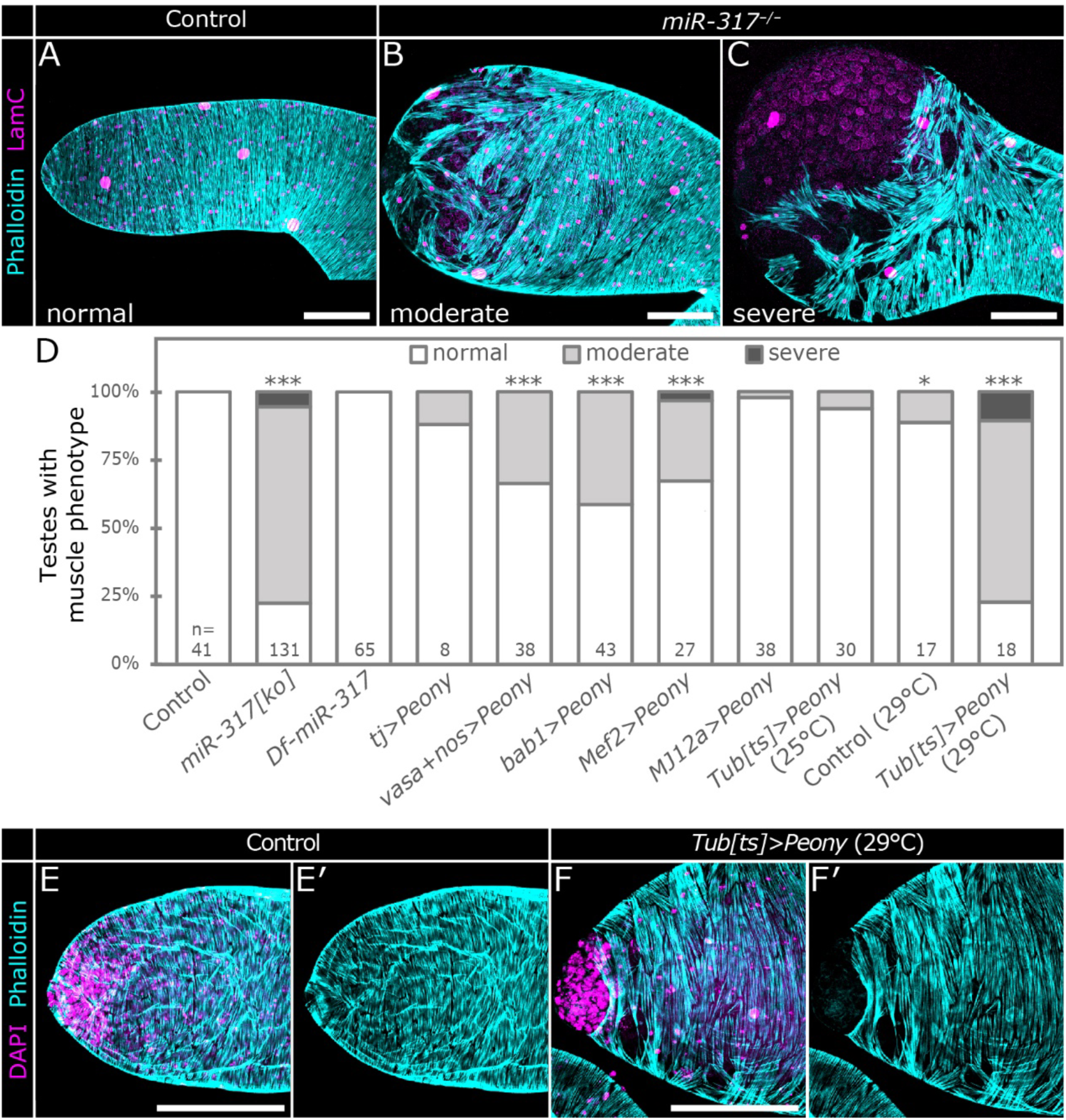
Regulation of the *Peony-miR-317* locus. **A** Control testis apex stained with Phalloidin (cyan) to mark the muscle sheath and anti-LamC (magenta) to mark nuclear envelopes. Multinucleated muscle cells have many small nuclei; pigment cells have large, sparsely spaced nuclei. **B-C** *miR-317^-/-^* testis apexes with moderate (B) and severe (C) muscle sheath disruption. In C the muscle sheath is severely disrupted, and the underlying germline is revealed by LamC staining. **D** Quantification of muscle disruption phenotypes in testes of various genotypes: *miR-317* mutants and *Peony* overexpression using cell- and tissue-specific drivers. **E** Apex of control testis stained for DAPI and phalloidin. Panel E′ depicts only phalloidin, showing an intact muscle sheath covering the apex. **F** Enlarged apex of testis overexpressing *Peony* under the control of *Tub-Gal80; Tub-Gal80[ts]* (*Tub[ts]>Peony*). Phalloidin stain (shown alone in panel F′) shows a broken muscle sheath at the apex. Scale bars in A, B, C, and E-F: 100 µm. In D: *, p<0.05; ***, p<10^-3^ as determined by chi-square analysis.

**Supplemental Figure 3.**
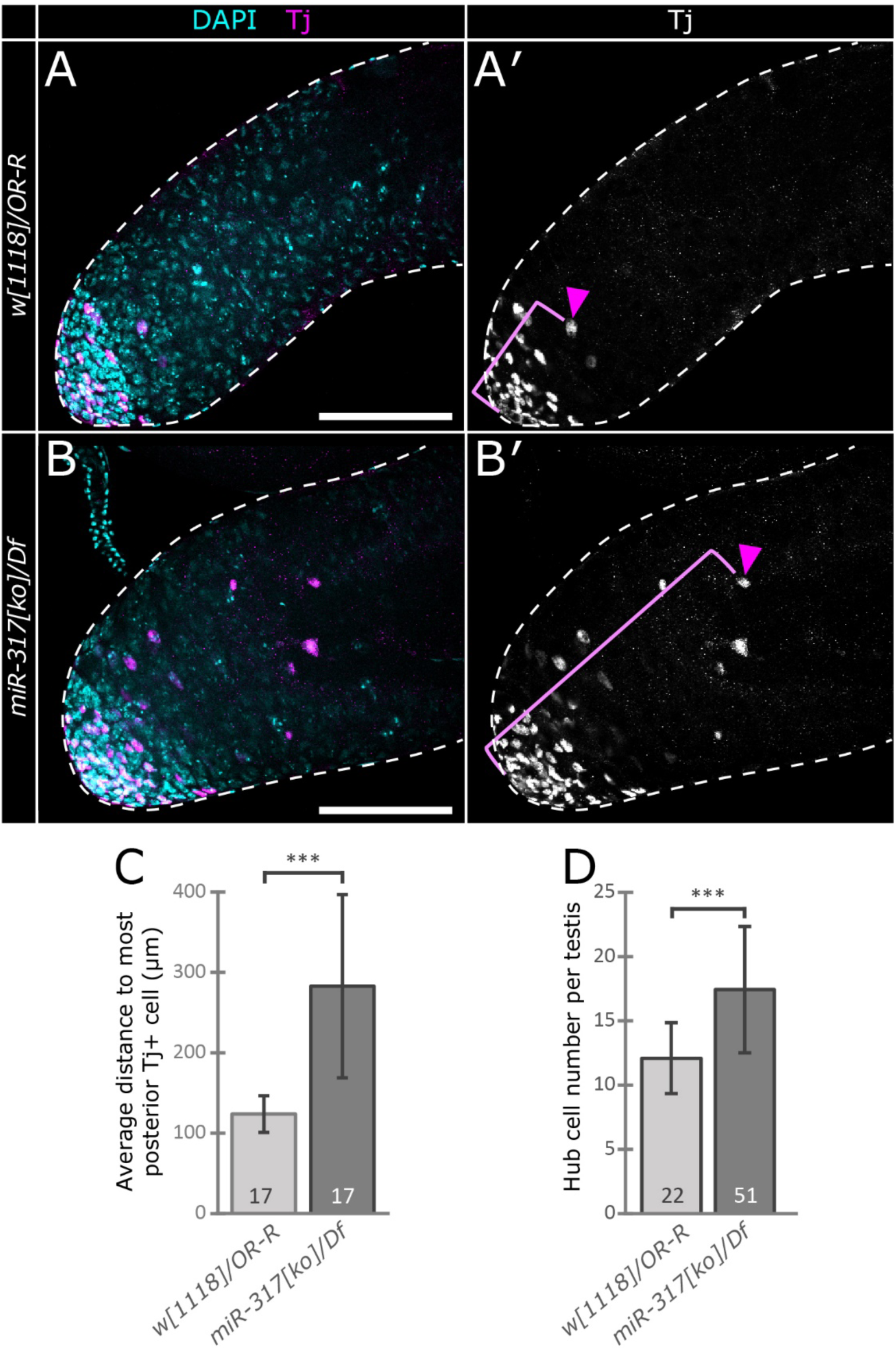
Effects of *miR-317* on testis somatic cells. **A-A**′ Control (*w[1118]/OR-R*) testis apical region in which Tj^+^ cells are clustered near the apex, reflecting the expression of Tj in hub cells, cyst stem cells, and early somatic cyst cells. **B-B**′ *miR-317[ko]/Df* testis apex exhibiting Tj^+^ cells clustered at the apex as well as a small number of Tj+ cells located more posterior to the DAPI-bright spermatogonia. In A′ and B′, Tj is shown in grayscale. Magenta arrowheads indicate the most-distal Tj^+^ cell, and brackets indicate distance from the apex. **C** Quantification of the average distance in µm from apex to most distal (posterior) Tj^+^ cell. **D** Quantification of the average number of hub cells per testis in controls (*w[1118]/OR-R*) and mutants (*miR-317[ko/Df]*). Error bars indicate standard deviation from the mean; Student’s unpaired t-test was utilized to test for statistical difference between genotypes based on number of testes (C) or hubs (D) quantified; ***, p<.001. Sample sizes given on bar graphs in C and D. Scale bars: 100 µm.

**Supplemental Figure 4.**
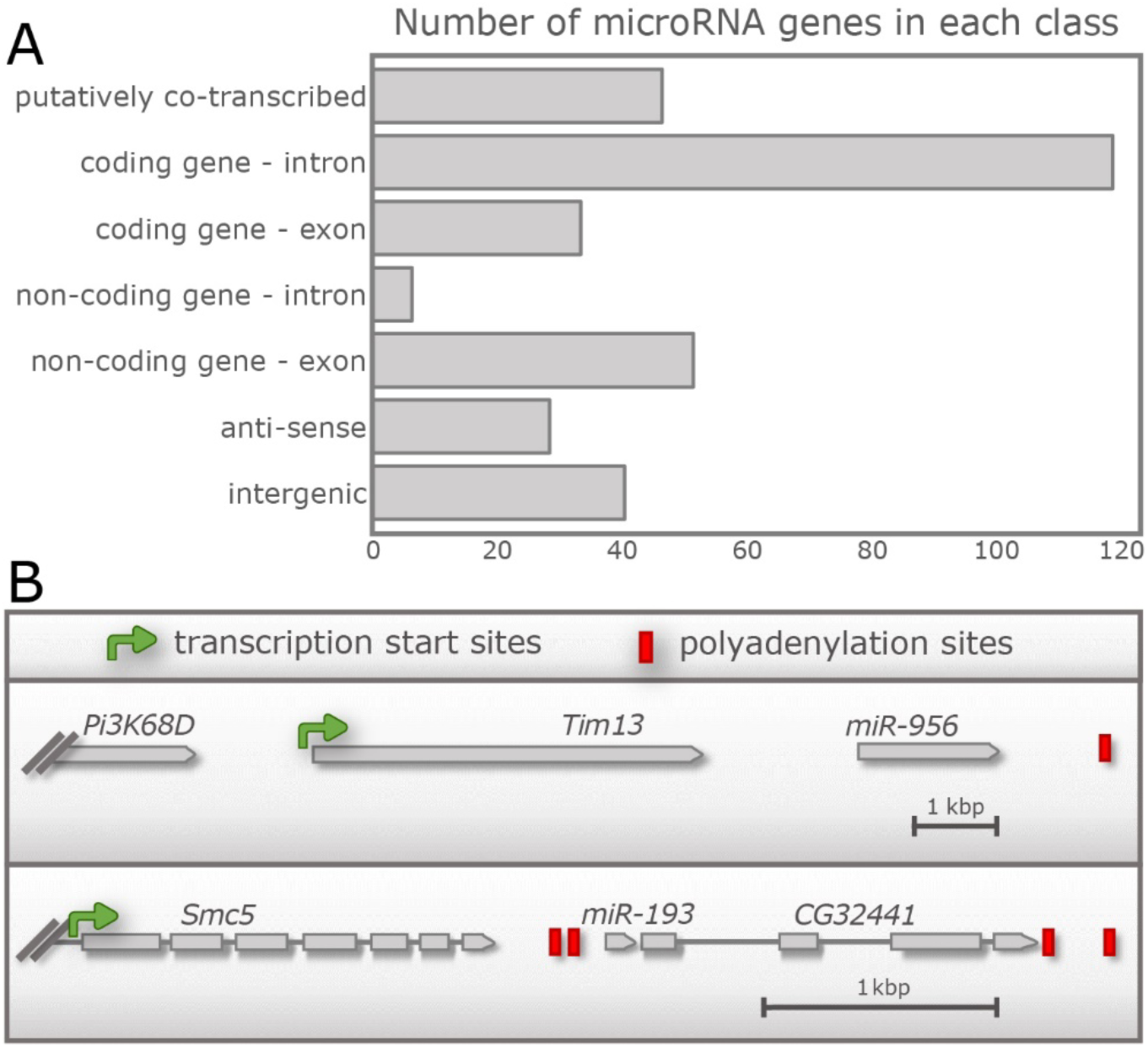
Putatively co-transcribed gene pairs in Drosophila. **A** Graph showing the number of Drosophila miRNA genes in each of several major categories. Putatively co-transcribed miRNAs reside closely upstream or downstream of other genes in the same orientation. **B** Examples of two putatively co-transcribed miRNA genes. *miR-956* is downstream of *Tim13*, and *miR-193* is immediately upstream of *CG32441*.

## Notes

### Competing Interest Statement

The authors have declared no competing interest.

### Summary of Updates

We added some new experiments and modified manuscript text.

